# A new heme enzyme family forms hydrazine groups in diverse biosynthetic pathways

**DOI:** 10.1101/2025.05.27.656397

**Authors:** Grace E. Kenney, Kwo-Kwang A. Wang, Tai L. Ng, Wilfred A. van der Donk, Emily P. Balskus

**Author notes:** Correspondence and requests for materials should be addressed to E.P.B. G.E.K is currently at the Department of Chemistry, University of Pittsburgh, Pittsburgh PA 15260, USA. K.-K.A.W. is currently at Bullseye Biosciences, Allston 02134, MA, USA.

## Abstract

Nitrogen-nitrogen (N–N) bond formation is an inherently challenging chemical process that plays a key role in the global nitrogen cycle. An array of microbial metalloenzyme complexes has evolved to shuttle nitrogen between biologically accessible reduced or oxidized states and its inert form as dinitrogen (N_2_) gas. More recently, N–N bond formation has been observed in a more specialized context, natural product biosynthesis. Here, we report the discovery of a unique metalloenzyme complex that forms hydrazine functional groups in the biosynthetic pathways of structurally diverse natural products. This heterodimeric system consists of a heme enzyme from a previously unidentified family and a partner ferredoxin.

Together, these enzymes effect the unprecedented four-electron reduction of nitrite (NO_2_^−^) to form a hydrazine functional group on a substrate amino acid in an oxygen-independent reaction that resembles primary microbial nitrogen metabolism. These enzymes are unexpectedly widespread among bacteria and are present in diverse genomic contexts, including cryptic biosynthetic gene clusters, highlighting the importance of this previously uncharacterized protein family.

TOC

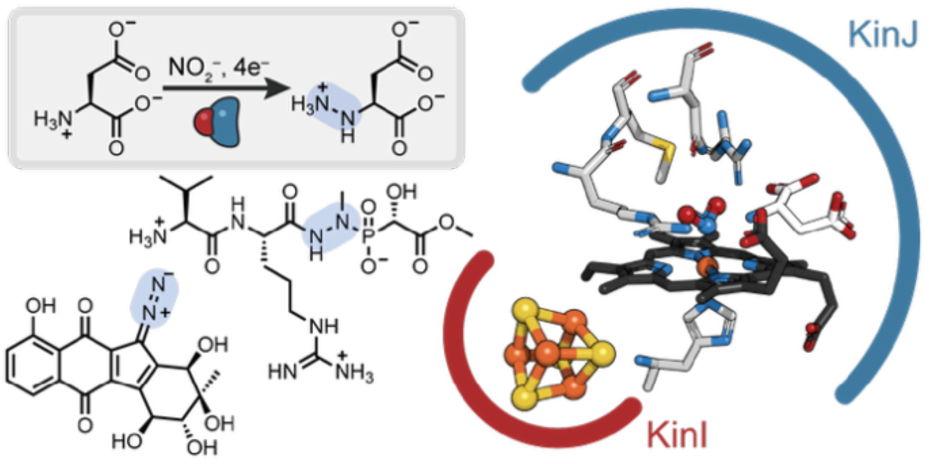

## INTRODUCTION

The Earth’s nitrogen cycle is defined by the formation and cleavage of N–N and nitrogen– oxygen (N–O) bonds.^1^ The predominant form of nitrogen on the planet is gaseous N_2_, but living organisms must either reduce this inert gas to ammonia (NH_4_^+^)^2^ or oxidize it via several enzymatic steps to form nitrite (NO_2_^−^) or nitrate (NO_3_^−^)^3^ in order to render it bioavailable. N_2_ can also be biologically regenerated, and microbes play central roles in every step of this process.^4^ In the nitrogen cycle, two distinct pathways forge N–N bonds, an intrinsically challenging chemical reaction. In the denitrification pathway,^5^ NO_3_^−^ is reduced to NO_2_^−^ by periplasmic or cytoplasmic nitrate reductases; NO_2_^−^ is then reduced to nitric oxide (^•^NO) via any of several unrelated metalloenzyme families that act as nitrite reductases.

Finally, two ^•^NO molecules combine with the aid of nitric oxide reductase to form nitrous oxide (N_2_O), which is later reduced to N_2_ by nitrous oxide reductase. In the anaerobic ammonium oxidation (anammox) pathway,^3^ NO_2_^−^ is reduced by nitrite reductases to form NO, and hydrazine synthase catalyzes N–N bond formation between ^•^NO and NH_4_^+^, yielding hydrazine (N_2_H_4_), which is ultimately oxidized by hydrazine dehydrogenase to form N_2_. While some components of the nitrogen cycle require oxygen, both the denitrification and anammox pathways are anaerobic, and metalloenzymes catalyze all steps in both pathways.

Enzymatic N–N bond formation is not limited to the nitrogen cycle. In recent years, a surprising range of enzyme families have been implicated in N–N bond formation in microbial natural product biosynthesis.^6–15^ These enzymes use diverse cofactors and their reactions proceed through disparate chemical mechanisms, but all share one notable feature: an oxygen-dependent step is required to activate one of the two nitrogen-containing reaction partners prior to N–N bond formation. Despite this progress, the enzymes responsible for constructing N–N bond-containing functional groups in many natural products are still unidentified, including fosfazinomycin, a phosphonate natural product^16,17^ that contains an internal hydrazide group, and the kinamycins and lomaiviticins, diazobenzofluorene polyketides (**Fig. 1a**).^18–20^ Identification of the biosynthetic gene clusters for these structurally unrelated natural products unexpectedly revealed that the *fzm*,^21^ *kin*,^22^ and *lom*^23,24^ pathways share several genes (**Fig. 1b**). Previously, we demonstrated via a combination of feeding studies, *in vivo* experiments, and *in vitro* biochemical characterization that the enzymes encoded by these genes generate a hydrazine synthon carried by an external scaffold prior to introduction into the final natural products; further, these studies implicated the involvement of NO_2_^−^ for the formation of L-hydrazinosuccinate (L-Hzs) as an early intermediate (**Fig. 1c**).^25,26^ However, the enzyme responsible for the proposed N–N bond-forming reaction between L-aspartate (L-Asp) and NO_2_^−^, yielding the key intermediate L-Hzs, has remained experimentally unidentified.

**Fig. 1.**
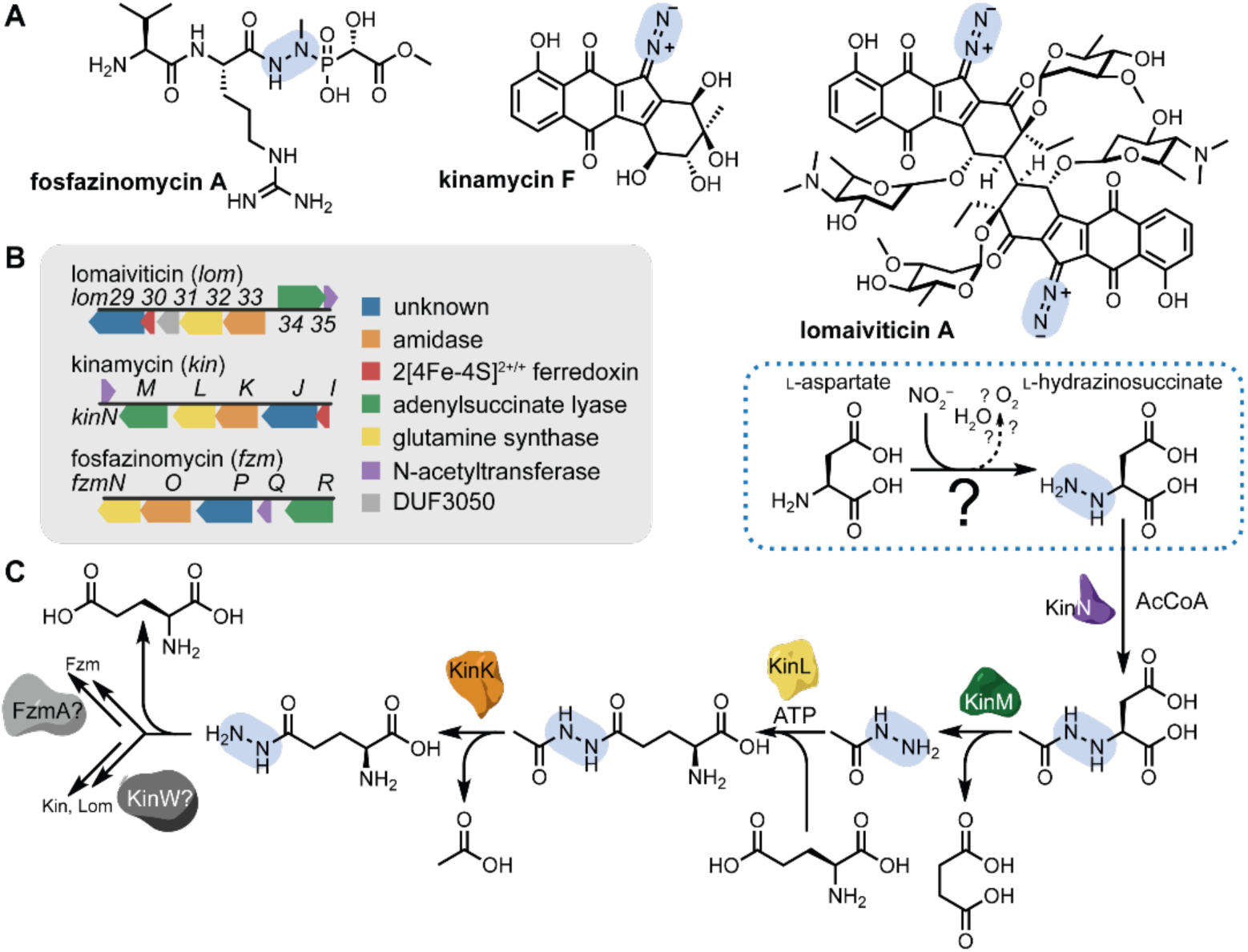
Natural products containing N–N bonds formed by a “hydrazino cassette.” **A**) Structures of fosfazinomycin, kinamycin, and lomaiviticin. **B**) The conserved hydrazino cassette of genes found in the *fzm*, *kin*, and *lom* biosynthetic gene clusters. **C**) The roles of most of the components of this pathway have been identified, but the role of the enzymes responsible for N–N bond formation (dashed box) remained purely hypothetical. The biosynthetic pathway splits off after the production of _L_-hydrazinoglutarate: for kinamycin and lomaiviticin, an *O*-methyltransferase-like enzyme installs the diazo group on the benzofluorene scaffold,^28^ while the subsequent pathway in fosfazinomycin has not yet been confirmed.

Here, we report the identification and characterization of the N–N bond forming metalloenzyme responsible for the generation of hydrazine groups in the biosyntheses of fosfazinomycin, kinamycin, and lomaiviticin. Unique among biosynthetic N–N bond forming enzymes, this heterodimeric complex—consisting of a previously unidentified class of heme enzyme and a ferredoxin partner—employs an oxygen-independent strategy, using NO_2_^−^ or NO_3_^−^ and an amine-containing substrate to form a hydrazine moiety in a net 4- or 6-electron reduction. Though this transformation resembles reactions involved in inorganic nitrogen assimilation, it is unique among all biological N–N bond forming reactions. This enzyme system is unexpectedly widespread and is proposed to be involved in the biosynthesis of both well-established and entirely new natural product families—a fact that is underscored by the recent report that a KinJ homolog is involved in the biosynthetic pathway for negamycin.^27^

## RESULTS

### KinJ and FzmP are members of a new heme enzyme family

At the outset of our studies, the most likely candidate for the proposed N–N bond forming reaction in these pathways was a hypothetical protein (FzmP, KinJ, and Lom29) encoded in the “hydrazino cassettes” of all three gene clusters that not yet have an assigned role (**Fig. 1b**). KinJ had no identifiable domains or cofactor-binding motifs, did not belong to a previously identified protein family, and its amino acid sequence provided no guidance to its potential structure, function, or enzymatic requirements. However, its proximity to a gene encoding a predicted ferredoxin (KinI, Lom30) in the *kin* and *lom* gene clusters was suggestive of possible participation in redox chemistry. To investigate the activity of this enzyme, recombinant KinJ from *Streptomyces murayamaensis* ATCC 21414^22^ as well as FzmP from *S.* sp. S-149 and *S.* sp. XY332 were heterologously expressed in *Escherichia coli* and purified via nickel affinity chromatography. Though no identifiable metal or cofactor binding motifs are present in the sequence of these proteins, the purified proteins are reddish-brown in color.

This unexpected observation led us to hypothesize that FzmP and KinJ both contain a heme cofactor. Supporting this proposal, spectral features consistent with the presence of a heme cofactor were identified via UV-vis spectrophotometry, and both the pyridine hemochromogen assay^29^ and LC-MS analysis of the protein and of the extracted heme confirmed the presence of a significant population of ferric heme *b* in aerobically expressed and purified protein (**Fig. 2A-B, S2A**). Although the heme loading levels in isolated enzymes were initially low, they could be increased to over 50% by either loading post-purification or by supplementing the growth media with iron and the heme metabolic precursor δ- aminolevulinic acid. In aggregate, these results support the identification of FzmP and KinJ as heme-containing proteins.

**Fig. 2.**
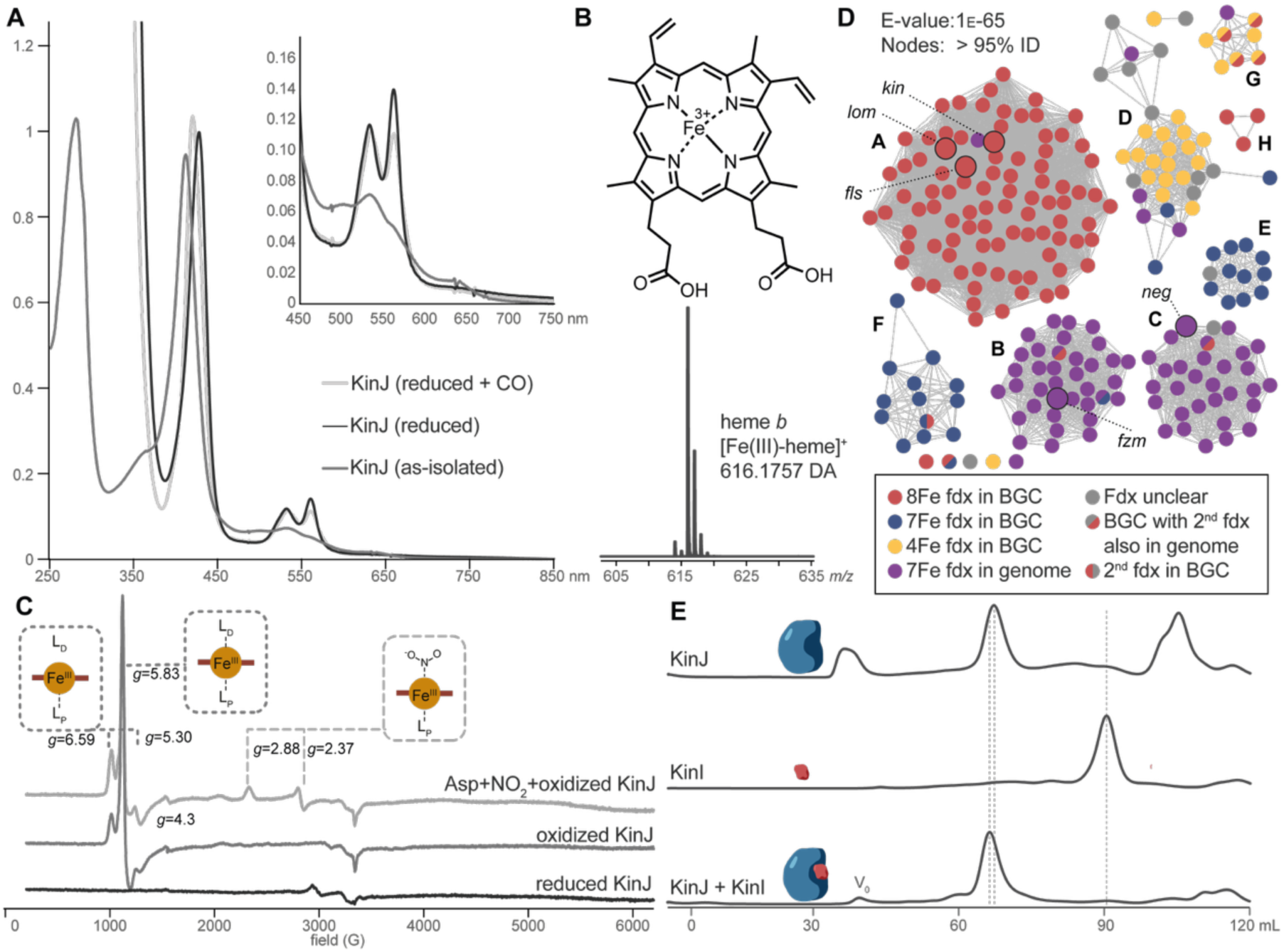
KinJ binds heme *b*. **A**) UV-vis spectrum of oxidized (red), reduced (blue), and reduced CO- bound (yellow) His_6_-KinJ. The Soret peak of the CO-bound species precludes P450-like behavior. **B**) Mass spectrum of extracted heme from His_6_-KinJ and heme *b* structure. **C**) EPR for KinJ shows two ferric high-spin species in as-isolated enzyme. Addition of NO_2_^−^ to the ferric enzyme results in the formation of a low-spin rhombic species. **D**) Analysis of the 10 genes to either side of *kinJ* homologs indicates that several distinct types of electron-transporting [4Fe-4S] ferredoxins are present in *kinJ* genomic neighborhoods. Superimposed on a sequence similarity network of KinJ homologs, it is clear that two subgroups of KinJ family members are found only in genomic contexts lacking electron transfer proteins, though – as will be discussed at greater length later – [4Fe-4S] ferredoxins are present elsewhere in many genomes. Highlighted representative nodes represent families of known function including KinJ from the kinamycin producer *S. ambofaciens* ATCC 23877, Lom29 from the lomaiviticin producer *Salinispora pacifica* DPJ-0019, FlsU2 from the fluostatin producer *Micromonspora rosario* SCSIO N160, FzmP from the fosfazinomycin producers *Streptomyces* sp. XY332 and *Streptomyces* sp. NRRL S-149, and NegJ from the negamycin producer *K. purpeofusca* NRRL B-1817.^27^ **E**) Anaerobic size exclusion chromatography indicates that KinI and KinJ form a heterodimeric complex when co-expressed.

Reduction of the KinJ and FzmP heme required sodium dithionite rather than ascorbate, indicating a low redox potential.^30^ Via UV-vis spectroscopy, we observed the core spectral features found in all hemes, including the Soret peak at 411 nm (427 nm in the reduced protein) and the α and β peaks associated with the Q-band at 532 and 560 nm in the reduced protein. Addition of carbon monoxide resulted in slight broadening of the Q-band features and a hypsochromic shift of the Soret peak to 420 nm; the lack of a 450 nm feature and of identifiable sequence motifs suggest that KinJ is not a member of the cytochrome P450 family (**Fig. 2A**). Electron paramagnetic resonance spectroscopy suggested that in oxidized KinJ and FzmP, the heme is predominantly in a mix of two high-spin configurations, namely a rhombic 5-coordinate environment and an axial six-coordinate environment (**Fig. 2C**),^31^ with the latter form predominant in FzmP. Absorption spectroscopy further supports these results: additional features were frequently observed above 630 nm when oxidized, and weak features in that region were also associated with high-spin hemes in other proteins.^32^ The reduced enzyme was EPR-silent.

Addition of L-aspartate did not alter the spectroscopic features of KinJ or its homologs, but addition of NO_2_^−^ (a strong field ligand) was associated with the formation of a 6-coordinate rhombic low-spin species (as well as a decrease in the populations of all other spin states and species) (**Fig. 2C**). NO_2_^−^ binding to heme *b* cofactors in other enzymes—including globins^33^ and nitrophorins^34^—results in the formation of species with g-tensor values in the same range. In nitrite reductases and nitrophorins, NO_2_^−^ binding to the cofactor is predominantly via nitrogen (*N*-nitro binding), while His-mediated H-bonding is thought to stabilize an alternate binding mode via oxygen (*O*-nitrito binding) in globins^34,35^. While the binding mode of NO_2_^−^ to KinJ is not yet clear, these results are consistent with the direct binding of NO_2_^−^ but not the proposed amino acid co-substrate to the heme.

With the identification of KinJ/FzmP as heme enzymes capable of binding NO_2_^−^, we next attempted to confirm their role in L-Hzs formation by incubating purified enzyme with NO_2_^−^ and L-aspartate (L-Asp). Though the enzyme was assayed under a wide range of conditions, including addition of various NTPs and other cofactors, metals, alternate substrates and nitrogen sources, alternate FzmP homologs, additional enzymes from the hydrazino cassette, spinach ferredoxin and its reductase, and cell lysate, no formation of L-Hzs was observed.

### KinJ homologs require a low-potential ferredoxin partner

This lack of activity led us to propose that a missing factor was required for N–N bond formation by KinJ/FzmP. The absence of identifiable sequence motifs made it difficult to envision possible additional cofactors or their binding sites in KinJ and FzmP. However, the unexplained presence of a ferredoxin in the *kin* and *lom* clusters prompted an in-depth analysis of the genomic neighborhoods of KinJ family members to identify potential partner proteins. Sequence similarity networks were constructed for an initial dataset of 869 KinJ homologs, the genomic neighborhoods (+/- 10 genes) of the 234 homologs chosen as representative nodes at 95% ID were examined, and hierarchical clustering was used to identify similar sets of genomic neighborhoods (**Fig. 2C**).^36,37^ We found that at this representative node level, 55% of KinJ gene neighborhoods (including the *kin*^22^ and *lom*^23,24^ gene clusters as well as gene clusters associated with the related benzofluorene compounds known as fluostatins (*fls*)^38^) include a gene predicted to encode a ferredoxin with one or two [3/4Fe-4S] clusters. Though the *fzm* gene cluster lacks such a gene,^21^ the apparent association between the greater family of heme-binding proteins and electron-transporting ferredoxin partners nevertheless suggested the latter might enable the redox chemistry performed by KinJ and its homologs.

To test this hypothesis, KinJ and FzmP were expressed alone or co-expressed with the ferredoxin KinI, encoded adjacent to *kinJ* in *S. murayamaensis*.^39^ Hexahistidine (His_6_) affinity tags were installed on one or both proteins at either the N- or C-terminus depending on the experiment, and proteins were initially expressed aerobically and purified anaerobically. KinI expressed alone with either N- or C-terminal His_6_ tags exhibited the spectral features and cofactor-binding characteristics associated with 2[4Fe-4S]^2+/+^ ferredoxins, albeit with evidence of partial cluster degradation (**Fig. S3A-D**).^30^ Gratifyingly, when co-expressed, both KinI and KinJ pulled each other down, regardless of tag location (**Fig S4A-C**), indicating the formation of a tightly bound complex. FzmP also co-purified with KinI, but only when the affinity tag of the ferredoxin was on the N-terminus (**Fig. S4B**). Size exclusion chromatography indicated that KinJ was monomeric when expressed alone, while co-expressed KinIJ formed heterodimers (**Fig. 2E**). FzmP and N-terminally His_6_-tagged KinI also formed heterodimeric complexes when co-expressed, but FzmP expressed along with C-terminally His_6_-tagged KinI formed a homodimer (**Fig S8B**). The strong association between the ferredoxin KinI and both heme proteins hinted that a heterodimeric complex may be critical for enzymatic activity.

### KinI and KinJ together form L-Hzs from L-Asp and NO_2_^−^

We next assayed KinJ and FzmP for activity in the presence of KinI. KinJ and FzmP were incubated with L-aspartate, NaNO_2_, KinI (either added separately or co-purified), sodium dithionite, and a low-potential electron mediator such as methyl or benzyl viologen; all assays were carried out anaerobically. After 3 kDa ultrafiltration to remove the proteins, the reactions were analyzed via liquid chromatography-mass spectrometry (LC-MS). A mass consistent with that of L-Hzs was observed, indicating the formation of an N–N bond-containing product from L-Asp and NO_2_^−^. This mass was observed only when all reaction components were present, and it shared a retention time, mass ([M-H]^−^ *m/z* 147.0404), and MS2 fragment ions with a synthetic L-Hzs standard (**Fig. 3A, S5, S6**). This result clearly identifies KinJ/FzmP as the key N–N bond forming enzyme in L-Hzs formation. It additionally establishes an explicit requirement for the binding of a ferredoxin partner: chemical reductants alone do not result in KinJ/FzmP activity.

**Fig. 3.**
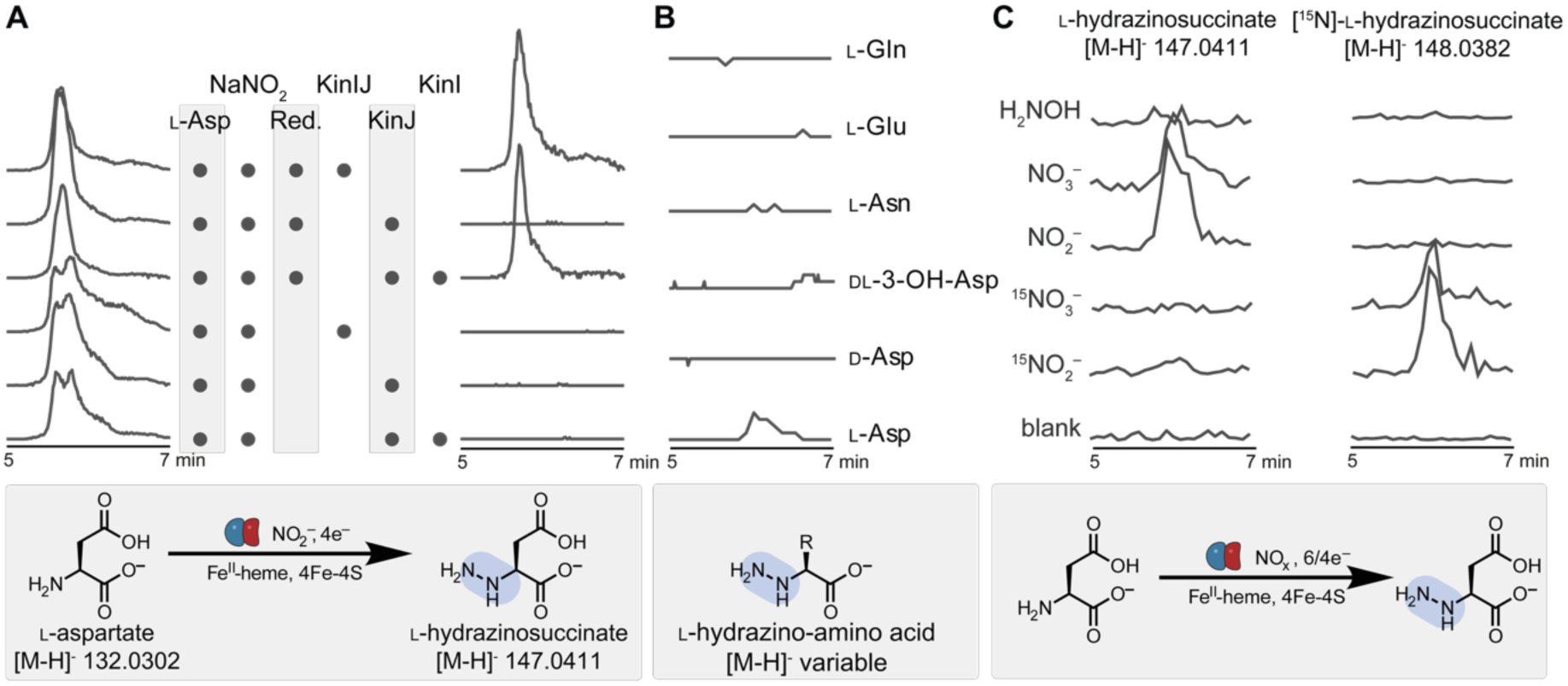
KinJ catalyzes _L_-hydrazinosuccinate synthesis from NO_2_^−^ and _L_-aspartate in the presence of KinI. **A**) Extracted ion chromatograms for _L_-aspartic acid and _L_-hydrazinosuccinic acid (shown scaled to the highest peak for that ion) confirm that _L_-hydrazinosuccinic acid is produced by KinJ in the presence of the _L_-aspartate and NO_2_^−^ co-substrates, a chemical reductant (“red.”, here sodium dithionite), an electron mediator (methyl or benzyl viologen), and the ferredoxin KinI. **B**) KinJ does not modify other amino acid substrates. Shown are the most structurally similar amino acids (_D_- Asp, _DL_-3OH-Asp, _L_-Asn, _L_-Glu, _L_-Gln), but no activity was observed with any other amino acid (**Fig. S7**). **C**) KinJ (and FzmP, not shown) do not produce _L_-Hzs when H_2_NOH is provided instead of NO_2_^−^. A decreased level of activity is seen when enzymes are provided with NO_3_^-^ rather than NO_2_^−^, but activity is observed.

**Fig. 4.**
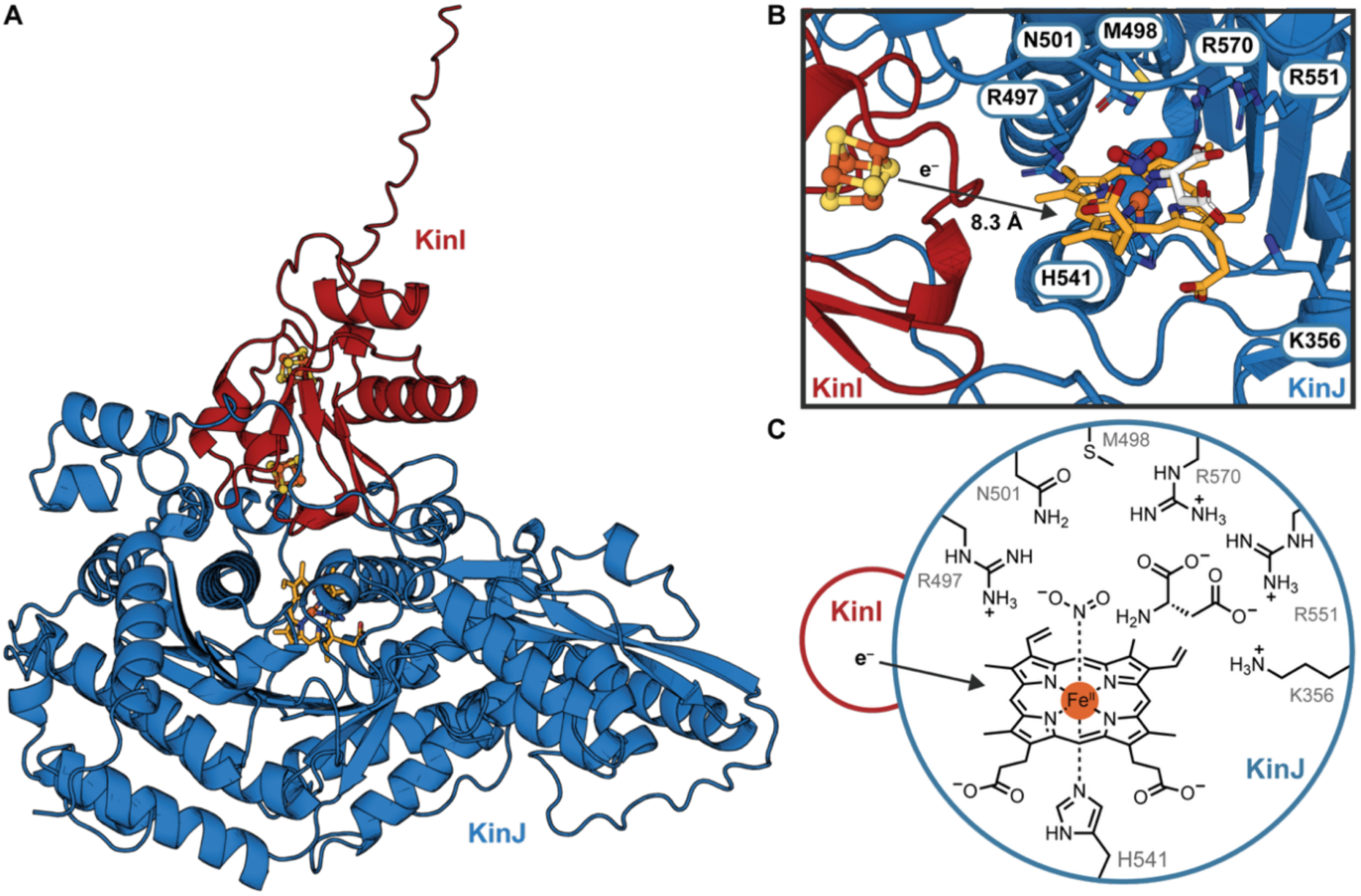
AlphaFold3 predicted structure for the KinIJ complex and a possible active site. **A)** Although KinJ has yet to be crystallized alone or in complex with KinI, detailed structural predictions computed by AlphaFold3 provide insight into a possible KinIJ complex (KinI in red, KinJ in blue). KinJ contains two domains, with the C-terminal domain better-conserved and with a distant similarity to GAF domain proteins; no clear structural precedents for the N-terminal domain have been identified. **B**) Proposed KinJ active site. The closer Fe-S cluster in KinI is within electron transfer distance; access to the heme binding pocket is potentially afforded by one of two predicted tunnels. **C**) Schematic of proposed KinJ active site, where a trio of conserved arginine residues may stabilize the negatively charged substrates. The presence of basic residues to coordinate and position the NO_2_^−^ substrate – and to act as proton donors – is a common feature in nitrite reductases.^53–55^

**Fig. 5.**
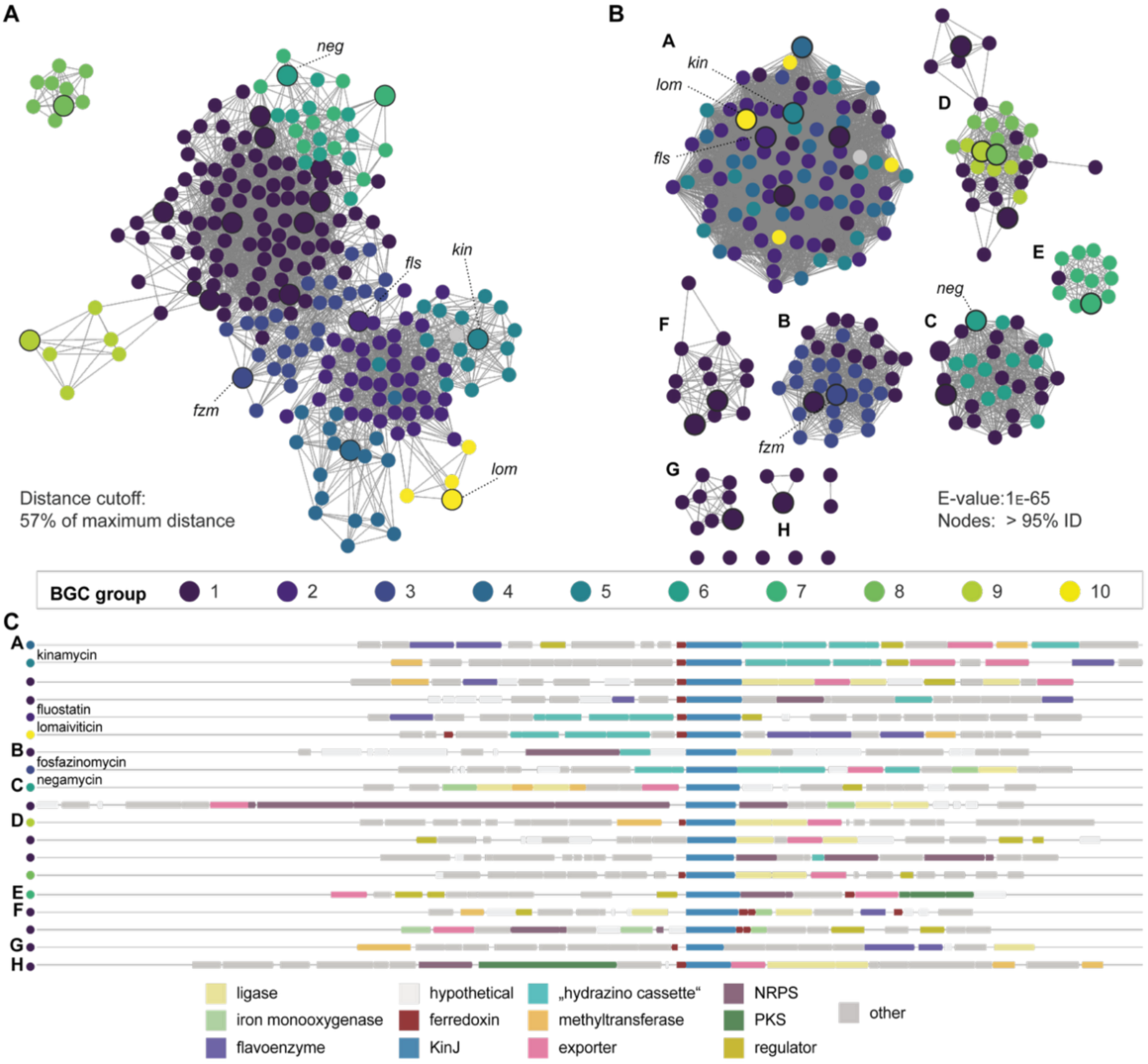
KinJ homologs are present in diverse biosynthetic pathways. **A**) In this genome neighborhood similarity network, nodes represent genome neighborhoods and edges between nodes represent the Euclidean distance between the *n-*dimensional vectors representing each genome neighborhood. To construct this network, edges with a Euclidean distance >57% of the maximum distance were trimmed, and clusters of nodes with similar connectivity are identified to highlight subsets of genome neighborhoods. Distinct clusters appear for kinamycin, fosfazinomycin, lomaiviticin, and fluostatin gene clusters, but a range of clearly divergent gene clusters are additionally present. **B**) When these genomic neighborhoods are mapped onto a sequence similarity network, where nodes represent KinJ homolog sequences and edges represent pairwise BLAST e- value scores better than a 1_E_-65 cutoff, it becomes clear that genome neighborhood type and sequence identity are not closely linked, indicating that closely related KinJ-like enzymes may be involved in divergent biosynthetic pathways. A subset of members of cluster A are in BGCs that encode diazobenzufluorenes, but many are embedded in entirely unrelated pathways. **C**) A sampling of genome neighborhoods chosen to maximize gene cluster diversity and KinJ sequence diversity illustrates the divergent pathways that involve KinJ homologs. All gene clusters include enzyme classes commonly associated with biosynthetic pathways, including ligases, methyl- and acetyl- transferases, iron and flavin monooxygenases, and dedicated exporters from the MFS superfamily.

**Figure 6.**
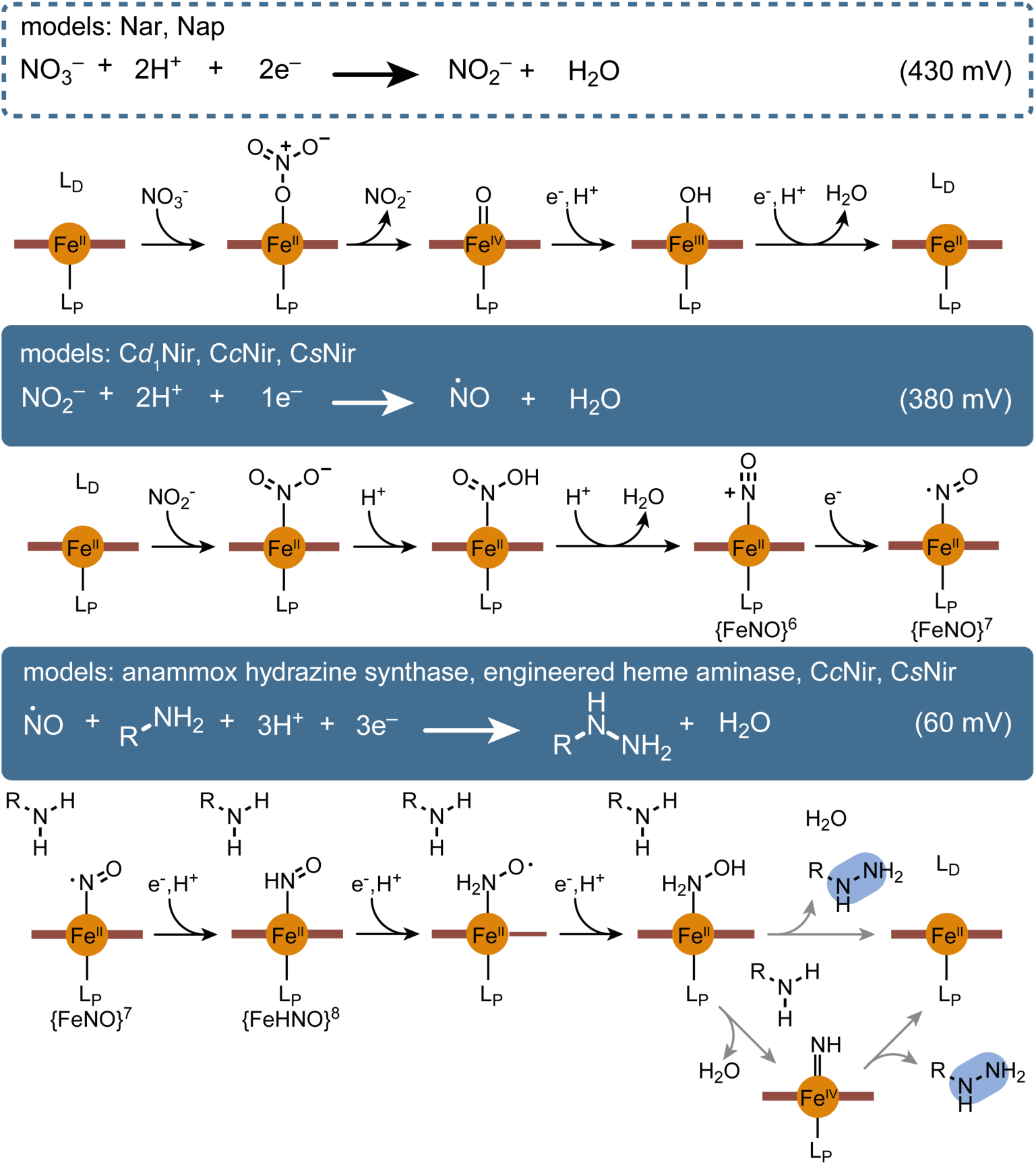
Enzymes providing possible models for production of L-Hzs by KinJ. Drawing analogies to known enzymes from the inorganic nitrogen cycle, NO_3_^−^ binds the heme active site and is reduced to NO_2_^−^ via a two-electron reduction; known respiratory and periplasmic nitrate reductases (Nar & Nap) use a different metallocofactor (MoCo),^77^ but if KinJ nitrate reduction follows a similar path, NO_2_^−^ might be released prior to further reduction. Next, NO_2_^−^ undergoes a single electron reduction to form ^•^NO, a reaction carried out by cytochrome *cd_1_* nitrite reductase (C*d*_1_Nir). Unlike that reaction, electrons are transferred via the partner ferredoxin, and the product ^•^NO is not released.^92^ Instead, an additional net 3 e^−^ reduction occurs, potentially via steps similar to those seen in ammonia- producing hexaheme cytochrome c nitrite reductase or siroheme/ferredoxin-dependent nitrite reductase (C*c*Nir and CsNir).^81,83^ In one possibility, the resulting hydroxylamine species engages in N–N bond formation with the target amino acid amine group. This is most similar to the proposed mechanism of anammox hydrazine synthase and is shown as the top path.^86,93,94^ This also has direct parallels to the widespread amine-hydroxylamine reactions seen in natural product N–N bond biosynthesis. However, there are other possibilities. A heme enzyme originally engineered to carry out CH amination using H_2_NOH via a nitrene intermediate was recently reported to also accept NO_2_^−^ as a nitrogen source,^89,95^ and N–N bond formation between nitrenes and amines has been observed or proposed in non- enzymatic contexts. This includes hydrazide formation mediated by organic N–O containing substrates and a catalytic Fe.^90,91^ It is conceivable that N–N bond formation in KinJ might also run through a nitrene intermediate (bottom path) en route to hydrazide formation. The nitrene is shown here as an Fe(IV) imido. While it can also be formulated as an Fe(III) imidyl radical or an Fe(II) nitrene adduct, the Fe(IV) imido has clear parallels to the Fe(IV) oxo species that is central to heme monooxygenase activity. Intriguingly, previous reports of heme-derived catalysts that generate nitrene intermediates suggest that a reduction potential of ∼-300 mV is associated with maximally effective nitrene transfer, and this appears to be consistent with the redox potential for KinJ.^96^ In either case, the initial NO_2_^−^ (or NO_3_^−^) reduction step or steps might make a low-redox potential ferredoxin partner to mediate electron transfer even more crucial.

Notably, homodimeric FzmP was unable to complete the reaction in the presence of added KinI, indicating that access to the heme enzyme by the ferredoxin is required, and that homodimer formation prevents this interaction (**Fig. S8A-D**). A requirement for co-expression and anaerobic co-purification with an appropriate ferredoxin partner may provide a rationale for the failure of early FzmP experiments to produce L-Hzs. Also importantly, the observed activity of both KinJ and FzmP is relatively low, with experiments in the presence of excess substrate and reductant rarely producing more than a single turnover and with high preparation-to-preparation variability. Efforts to chemically reconstitute KinI in the presence of KinJ or FzmP failed, complicating attempts to provide better-loaded ferredoxin partners to KinJ homologs. Finally, activity was only observed with the use of a low-redox-potential electron mediator (predominantly methyl or benzyl viologen) (**Fig. S7A**). In the absence of an electron mediator or in the presence of a higher potential electron mediator, activity was significantly diminished (phenosafranin) or absent (anthraquinone-2,6-disulphonate, Nile blue B, and 1-MeO-5-PMS). These observations suggest that additional, as-yet-unidentified factors may be limiting activity. Nevertheless, these data strongly support the hypothesis that catalysis of the net 4-electron reduction of NO_2_^−^/ L-Asp to L-Hzs by KinJ and FzmP requires a dedicated ferredoxin partner.

We next investigated the substrate promiscuity of both KinJ and FzmP. The twenty canonical L-amino acids, along with D-Asp and DL-3-OH-Asp, were examined for their reactivity with NO_2_^−^ as a reaction partner, but no substrate consumption or product formation was observed with any amino acid other than L-Asp (**Fig. 3B, S7B**). However, a recent report of a KinJ homolog in negamycin biosynthesis (NegJ) that produces hydrazine acetic acid from glycine indicates that some family members have distinct substrate preferences.^27^ Alternate nitrogen sources were also investigated. The reactivities of NO_3_^−^ and NH_2_OH were examined both alone and before or after incubation with NO_2_^−^ to examine the possibility of these molecules binding competitively to the enzyme (**Fig. 3C**). Diminished but replicable levels of activity were observed for KinJ when NO_3_^−^ was used as a substrate in lieu of NO_2_^−^, but significantly lower levels of NO_3_^−^-dependent activity were seen for FzmP. No activity was observed with NH_2_OH for either enzyme, although trace reactivity with NH_2_OH was reported for the recently published NegJ.^27^ In general, KinJ activity was less affected by pre-incubation with potential alternative substrates. This observation may imply that while the preferred KinJ reaction is the four-electron reduction of NO_2_^−^ to form the hydrazine group of L-Hzs, KinJ homologs can perform the equivalent six-electron reduction of NO_3_^−^ to form the same product, and that relative affinities for or reactivity with NO_2_^−^ and NO_3_^−^ may differ between homologs.

These experiments supported the identification of the KinJ/FzmP product, but did not indicate which nitrogen in L-Hzs was derived from NO_2_^−^. We thus compared reactions carried out using NO_2_^−^ and ^15^NO_2_^−^ with unlabeled L-Asp. In these reactions, L-Asp remains unlabeled, with ^15^N incorporation seen only in L-Hzs. Comparison of MS2 fragmentation of L- Hzs in the isotopically labeled and native-abundance products is consistent with localization of the isotopically labeled nitrogen to the distal position (**Fig. S9**). These results support a model where N–N bond formation occurs between NO_2_^−^ or a reduced nitrogen species and an intact amino acid co-substrate.

### Mutagenesis and structural predictions reveal a potential KinJ active site

To begin to decipher how the KinIJ complex forms L-Hzs, we sought to characterize the KinJ active site. Previously identified heme-binding motifs provided no guide for further identifying the heme and substrate coordination chemistry of KinJ. However, as hemes have a relatively limited repertoire of axial ligands, we sought to identify candidate heme-binding residues using bioinformatics. Highly conserved residues within the KinJ family include the most common axial ligand, histidine, but also the less frequently observed ligands methionine, cysteine, tyrosine, and lysine (**Fig. S10**). To help identify candidate axial ligands, a set of variants were produced in which these highly conserved residues were individually replaced with an alanine (or phenylalanine in the case of tyrosine).

Several variants exhibited altered heme *b* loading, as analyzed via UV-vis spectroscopy (**Fig. S11A**). Most strikingly, the H541A variant exhibited complete loss of heme binding, suggesting that His541 may be one of the axial ligands of the heme; a H541C variant restores heme binding, albeit with altered features consistent with a thiolate rather than an imidazole or imidazolate ligand (**Fig. S11D**).^40^ By contrast, the C468A variant (and to a lesser extent the M468A variant) exhibited a marked decrease in heme *b* content, and an increase in other heme species (evinced by mass spectrometry and spectral shifts, **Fig. S11B, C**). None of the variants exhibited enzymatic activity (**Fig. S11B**). Two variants with detectable heme exhibited shifted 5-coordinate high-spin or 6-coordinate low-spin NO_2_^−^- bound species (**Fig. S11C**). We note that loss of activity does not necessarily indicate that the residue replaced in a given variant is a heme ligand or within the active site; the highly conserved tryptophan and tyrosine residues might be involved in electron transfer from the ferredoxin. However, this analysis pinpoints a limited set of amino acids likely to be present in the proximity of the heme cofactor, including C468, M498, H504, and H541, with H541 as the most likely candidate for the axial ligand.

We next sought to identify a potential heme binding site in KinJ. As members of this enzyme family have yet to be structurally characterized, we turned to protein structure prediction.

AlphaFold2^41,42^ models of the apo KinIJ complex provided a potential rationale for the set of variants in which heme binding is altered. KinI and KinJ were modeled separately and together as a heterodimeric complex using AlphaFold2, with heme additionally added in AlphaFold3 models (**Fig. 4A-B, S12**).^43^ The tertiary structure and folds of KinJ are rendered similarly across protein structure prediction models with only small differences at the N- and C-termini and in a handful of labile loop regions (**Fig. S12**), with significant uncertainty (as indicated by pLDDT values) primarily predicted in those locations (**Fig. S12A,B**). H541, the candidate heme axial ligand, is located in a pocket in the C-terminal domain in close proximity to M498 and C468, which are two of the additional residues wherein alanine mutation perturbed the heme binding environment. Three highly conserved arginine residues positioned nearby (R497, R551, and R570) are well-positioned to bind and stabilize the two acidic substrates, NO_2_^−^ and L-aspartate (**Fig. 4B-C**). An equivalent binding location – with equivalent stabilizing interactions from active site residues – was modeled into the active site of the recently reported NegJ, and mutagenesis of that protein also supports identification of these key active site features.^27^

Intriguingly, the DALI and Foldseek servers^44,45^ identify distant structural homology in the C-terminal region of KinJ to two classes of sensor proteins containing GAF domains (**SI** **table 4**).^46,47^ In the first class, the GAF domain binds a single heme (with the mycobacterial sensor DosT^48^ as the closest identified hit), while in the second class, a set of bacteriophytochromes, the GAF domain binds heme-derived linear tetrapyrrole (bilin) chromophores (with cyanobacterial PixJ phototaxis regulators the closest identified hit).^49^ In both cases, the cofactors are located in regions equivalent to the proposed heme-binding site in KinJ (**Fig. S13A**). This may point towards an origin for the C-terminal domain of KinJ as a heme sensor; the origin of the N-terminal domain remains unclear.

AlphaFold3^43^ evocatively installs a heme cofactor in a roughly corresponding location, bound to H541 and with an open coordination site (**Fig. 4**). 4Fe-4S clusters can be modeled into the bound KinI partner using homology alignment against the closest experimentally studied ferredoxin, a 7Fe-8S variant of the 2[4Fe-4S]^2+/+^ *Allochromatium vinosum* “Alvin” ferredoxin (PDB: 3EUN).^50^ The edge-to-edge distance between the heme cofactor and the closest 4Fe- 4S cluster is 8.3 Å, well within cofactor distances experimentally determined to be compatible with direct electron transfer.^51^ The pocket above this location is sufficiently large to accommodate both a smaller ligand—such as NO_2_^−^ or NO_3_^−^ —in the open coordinate site and a larger ligand—such as an amino acid—immediately adjacent (**Fig. S13B**). This pocket may be accessible via at least one tunnel with a diameter of at least 1.9 Å (**Fig. S13C**).^52^ The area surrounding these potential ligand binding sites includes several highly conserved residues, including R570, R550, N501, R496, and potentially K356, all of which are positioned to support potential ionic or hydrogen bonding to negatively charged substrates (**Fig. 4B,C, S13B**). When combined with the biochemical characterization of KinJ variants, the structural models are consistent with the identification of a proposed heme binding site, with the iron-sulfur cluster of the partner ferredoxin located sufficiently closely to facilitate electron transfer.

### KinJ homologs are present in diverse biosynthetic pathways

While KinJ homologs share core sequence features and can thus be predicted to share major biochemical features such as heme binding, the genomic neighborhoods in which they occur are diverse. Genera that encode KinJ homologs are predominantly aerobic Actinomycetes (including *Bifidobacterium*, *Streptomyces*, *Corynebacterium*, *Micromonospora*, and *Nocardia*), but include some Firmicutes (*Staphylococcus, Streptococcus*), and even Gram-negative Proteobacteria (*Burkholderia*). It is noteworthy that most gene clusters containing KinJ homologs (56% using 95% identity representative nodes) are not predicted to produce phosphonate or diazobenzofluorene natural products (**Fig. 5**).

Indeed, additional components of the diazo cassette are typically absent, with most gene clusters encoding only a KinJ homolog and a partner ferredoxin along with other predicted biosynthetic enzymes (**Fig. 5C**).

Some subsets of gene clusters are associated with non-ribosomal peptide synthetase or polyketide synthase modules. Components of the “hydrazino cassette” other than KinI and KinJ are not conserved beyond the fosfazinomycin gene cluster and the gene clusters encoding (diazo)benzofluorene natural products, including fluostatin, kinamycin, and lomaiviticin. However, some features from other pathways with hydrazine intermediates are present: in cluster E, for example, BGCs encode homologs (>55% identity) of the Haa-succinylating enzymes Tri31/AzaB/NngQ and the succinyl-Haa adenylating enzymes Tri29/AzaC/NngO, which load succinyl-Haa onto a dedicated carrier protein for further modification^15,56,57^ This potentially implicates glycine as the substrate for KinJ homologs within that cluster, just as aspartate may be the substrate for members of clusters A and B, and glycine may be the substrate for members of cluster C.

Attempts to connect cryptic KinJ-encoding gene clusters to previously discovered bacterial natural products provided further support for the involvement of this enzyme family in N–N bond formation. One homolog was identified in the actinomycete *Kitasatospora purpeofusca* which produces negamycin, a hydrazide-containing antibiotic whose biosynthesis was uncharacterized at the time of this work.^58^ The *K. purpeofusca* genome encodes not just a KinJ homolog but enzymes involved in *N*-methylation, lysine hydroxylation, and formation of beta-lysine, all of which are colocalized and have clear potential roles in the production of negamycin (**Fig. 5C**). The recent report describing the discovery of NegJ supports the correct identification of this gene cluster.^27^ A preponderance of additional gene clusters encodes members of the ATP-grasp superfamily (including but not limited to the *neg* and *fzm* gene clusters), which are frequently involved in amide bond formation, suggesting that many pathways that use KinJ homologs are likely to produce peptidic or peptide-like compounds. However, some subsets are associated with PKS or NRPS modules, or with unidentified hypothetical proteins. Others are associated with genes encoding enzymes from azaserine, triacsin, and *N*-nitroglycine biosynthesis, which involves a distinct N–N bond forming enzyme,^15,56,57,59^ raising the intriguing possibility that KinJ and AzaEFG may be interchangeable sources of hydrazino amino acids in biosynthetic pathways. The diversity of these cryptic gene clusters suggests KinJ homologs are involved in the production of a wide array of yet-to-be discovered natural products containing N–N bonds, including not only hydrazide or diazo groups but potentially other nitrogen-rich structural motifs.

Additional features of these gene clusters may shed light on the reactions mediated by KinJ homologs. The use of NO_2_^−^ as a co-substrate for N–N bond formation in the *kin* and *fzm* systems was inferred from the presence of genes encoding CreDE homologs.^60^ These enzymes generate NO_2_^−^ *in situ* from L-aspartate in the biosynthesis of cremeomycin and other natural products. However, genes encoding CreDE homologs are absent from most gene clusters and genomes encoding KinJ homologs, suggesting that NO_2_^−^ (or perhaps NO_3_^−^) is in many cases produced or acquired via other means. Finally, while most gene clusters encoding KinJ family members appear to also encode a dedicated ferredoxin, the ferredoxins do not all belong to the same subtype and indeed are unexpectedly diverse (**Fig. S14**). KinI-like ferredoxins are low-potential 2[4Fe-4S]^2+/+^ (“8Fe”) proteins with an N-terminal extension that do not fit into the existing “Alvin” and “Clostridial” classes, while the ferredoxins encoded in gene clusters from *Burkholderia* and related organisms are low-potential “7Fe” ferredoxins with one [3Fe-4S] and one [4Fe-4S] iron-sulfur cluster.^61^ Several subgroups of KinJ homologs, including both FzmP and NegJ, frequently lack a ferredoxin encoded in their gene cluster. However, 7Fe ferredoxins similar to those in *Burkholderia* BGCs are encoded in >80% of these species’ genomes, raising the possibility that they may be able to substitute for a dedicated KinI homolog. Indeed, the study of the negamycin- producing enzyme NegJ utilized the 7Fe ferredoxin identified in the *K. purpeofusca* genome that is encoded outside of the negamycin BGC.^27^ *Staphylococcus* gene clusters encode a smaller ferredoxin with a single [4Fe-4S] cluster.^62^ Unexpectedly, while gene cluster components are otherwise not highly correlated with KinJ sequence similarity, the presence or absence of an in-BGC ferredoxin partner *is* correlated with specific subsets of KinJ homologs with similar sequences (**Fig. 2D**), possibly supporting specificity for ferredoxin partners in many systems. The relative positioning of the ferredoxin when binding the KinJ homolog is consistent in all KinJ-associated ferredoxins, with a predicted basic patch surrounding the tunnel to the KinJ active site mediating binding by a complementary acidic patch on the partner ferredoxin (**Fig. S15**). Additional work will clearly be necessary to fully understand the substrate scope of the KinJ family and its ferredoxin partners, as well as the diversity of N–N bonds present in the final natural products produced by these systems.

## Conclusions

KinIJ and NegJ, which was reported during the preparation of this manuscript, are the first characterized representatives of a new metalloenzyme family consisting of a heme enzyme and a partner ferredoxin.^27^ The data provided here, together with previous knowledge of fosfazinomycin and kinamycin biosynthetic logic, show that KinJ and its homolog FzmP catalyze the formation of L-Hzs from NO_2_^−^ and L-Asp. Certain aspects of the KinJ system resemble other N–N bond-forming enzymes in biosynthetic pathways. Several systems rely on NO_2_^−^, as KinJ does, but form N–N bonds via enzymes that are not currently thought to directly catalyze redox reactions including homologs of CreM in the cremeomycin pathway,^10,15,63^ a membrane protein AzpL involved in alazopeptin biosynthesis,^64^ and possibly the enzymes involved in alanosine biosynthesis, which may also use non-dedicated nitrite and nitrate reductases from the inorganic nitrogen cycle.^65–67^

A second and larger class of N–N bond forming enzymatic systems rely on the oxygen-dependent hydroxylation of an amine group, forming a hydroxylamine species that facilitates N–N bond formation. This includes a trienzyme system involved in the biosynthesis of the hydrazone-containing compound s56-p1 via a hydrazinoacetate intermediate, now identified in pathways for many additional compounds.^13,15,56,57,59,68–71^ *N*-hydroxylation is also seen in dixiamycin biosynthesis (XiaK)^7^ and in the biosyntheses of kutzneride (KtzIT) and other piperazic acid-containing compounds.^6,72,73^ Additional, more complicated oxidative pathways to N–N bond formation have been reported in the biosyntheses of streptozotocin (SznF), gramibactin, and chalkophomycin,^12,74,75^ as well as 8-aza-guanine (PtnF)^14^ and azoxymycin (AzoC).^11^ An additional new oxidase family, ToxC, has been proposed to facilitate N–N bond formation in toxoflavin biosynthesis.^76^ While there are many subtle mechanistic differences, and though the biosynthetic pathways of many N–N bond-containing natural products remain unresolved, all the pathways characterized to date involve an O_2_-dependent *N*-hydroxylation step prior to N–N bond formation or a non-redox incorporation of NO_2_^−^.

Unlike all other N–N bond forming biosynthetic enzymes, KinIJ catalyzes a comproportionation reaction that does not require O_2_, forming an N–N bond between NO_2_^−^ and the amino group of an amino acid to produce a hydrazine linkage in a redox-dependent and reductive process using a yet-to-be-identified NO_x_ intermediate. This unique reactivity resembles enzymes involved in the inorganic nitrogen cycle, particularly cytochrome *cd_1_* nitrite reductase and hydrazine synthase, suggesting these enzymes may be more relevant for understanding the mechanism of the KinIJ system. The KinIJ-catalyzed production of L-Hzs appears to be an O_2_-independent 4e^−^ reduction of NO_2_^−^ and L-Asp. Like inorganic NO_3_^−^ and NO_2_^−^–reducing systems, KinJ requires an external reductant and a partner domain or protein capable of electron transport.

The apparent nitrate reductase activity of the KinIJ system has one potential biological model in nitrate reductase; these membrane-bound (Nar) or periplasmic (Nap) enzymes are members of the DMSO reductase superfamily and the quinol-dependent 2e^−^ reduction of NO_3_^−^ to NO_2_^−^ occurs at their characteristic molybdenum pyranopterin active site, with electron transfer via a [4Fe-4S] site.^77^ However, Moco enzymes catalyze concerted 2e^−^ reduction reactions, while a single heme cofactor is likely to catalyze distinct 1e^−^ reduction steps, raising intriguing questions about the mechanism of nitrate reduction by KinIJ.

Cytochrome *cd_1_* nitrite reductase (C*d*_1_Nir)^78^ provides a possible model for an initial NO_2_^−^ reduction step, prior to N–N bond formation. In this enzyme, NO_2_^−^ binds a unique heme *d_1_* cofactor; electrons are channeled towards this active site via an additional heme *c* site, and NO_2_^−^ is reduced to form ^•^NO in a 1e^−^ reductive step. The nitrite dismutase activity of nitrophorins provides an alternate, if mechanistically complex, model for ^•^NO generation by NO_2_^−^ reduction.^79^ A recent review of the role of ^•^NO in natural product biosynthesis highlighted its involvement in N–N bond formation in the inorganic nitrogen cycle and the surprising dearth of biosynthetic enzymes taking advantage of this chemistry;^80^ the involvement of an ^•^NO intermediate in KinIJ-mediated N–N bond biosynthesis would fill that gap. Two other heme-dependent nitrite reductases may provide relevant models for subsequent reductive steps: the pentaheme cytochrome c nitrite reductase (C*c*Nir)^81,82^ and the ferredoxin-dependent siroheme-utilizing nitrite reductase (*C*sNir)^83^ – though there are limits to the similarity of these systems, both of which produce ammonia and rely on a complex network of metal centers. While there are other NO_2_^−^-reducing enzymes, systems that differ in both cofactor and end product are less likely to provide a relevant model for KinJ.^84,85^

When considering a route towards N–N bond formation, the most relevant enzyme to KinIJ in the inorganic nitrogen cycle is a unique hydrazine synthase (HZS) complex found in anammox bacteria. This enzyme utilizes a multiheme system to affect the comproportionation of hydrazine from hydroxylamine and ammonia.^86^ However, the NO_x_ species that is the initial substrate of HZS is ^•^NO, not NO_2_^−^ or NO_3_^−^: the 3e^−^ reduction of ^•^NO to hydroxylamine is accomplished at one heme site with electron transfer (and possibly ^•^NO retention) with the help of a partner protein,^87,88^ and the resulting hydroxylamine is channeled to a second heme site where the comproportionation reaction occurs. Uniquely, KinJ appears to combine the activity of hydrazine synthetase with that of a nitrite or even a nitrate reductase, and to do so at a single active site. This reaction is unprecedented in synthetic chemistry. However, one intriguing parallel is found in a heme-containing protoglobin that was originally engineered to use hydroxylamine to generate a nitrene group for CH-amination but that can also accept NO_2_^−^ as a precursor,^89^ raising the possibility the N–N bond formation by KinJ may also involve a nitrene intermediate. The piperazate-forming heme enzyme KtzT was also recently proposed to employ a nitrene intermediate in intramolecular N–N bond formation between the backbone and sidechain amines of the hydroxy-Lys substrate,^73^ and nitrene-mediated hydrazine and hydrazide formation is also known in synthetic chemistry.^90,91^

By analogy to these existing enzymatic systems, we hypothesize KinIJ may first catalyze the 1e^−^ reduction of NO_2_^−^ (or, with an additional 2e^−^ reduction step, NO_3_^−^) to ^•^NO (**Fig. 6, top & center**). At that point, it is conceivable that a further 3e^−^ reduction to hydroxylamine may occur before an N–N bond is formed directly between the heme-bound hydroxylamine group and the primary amine of the amino acid substrate, following the mechanistic logic common in other N–N bond-forming enzymes (**Fig. 6, bottom**). Alternately, the hydroxylamine intermediate might undergo water loss, generating a nitrene poised to from an N–N bond with the nearby amino acid primary amine (**Fig. 6, bottom**). It is also conceivable that the NO intermediate might be capable of N–N bond formation as part of a radical rearrangement, with further reduction and release occurring subsequently. All routes are consistent with isotopic labeling studies that indicate that the nitrogen derived from NO_2_^−^ is the distal nitrogen in L-Hzs. More work will be required to unravel the detailed mechanism for this enzyme family.

The redox-dependent yet oxygen-independent mechanism of N–N bond biosynthesis in the Kin, Lom, Fzm, and Neg systems is highly unusual, and KinJ homologs belong to a previously unknown and uncharacterized protein family. Elucidating the mechanism of this enzyme family will enhance our understanding of heme-nitrogen biochemistry, with implications that go well beyond natural product biosynthesis. Moreover, KinJ homologs are present in an unexpectedly large and diverse number of natural product biosynthetic pathways. These enzymes are found not only in Actinomycetes but also in Gram-negative bacteria, and perhaps most notably in human-associated bacteria, including both commensal genera (*Bifidobacterium*) and opportunistic pathogens (*Burkholderia*, *Nocardia*, *Staphylococcus*). The identification of KinJ’s reactivity now sets the stage for the discovery investigation of wide variety of novel natural products containing N–N bonds.

## Author Contributions

Grace Kenney: formal analysis, data curation, investigation, methodology, conceptualization, visualization, writing – original draft and review & editing. Abraham Wang: investigation, methodology, conceptualization, writing – review and editing. Tai Ng: investigation, methodology, writing – review and editing. Wilfred van der Donk: project administration, supervision, funding acquisition, methodology, conceptualization, writing – review & editing. Emily Balskus: project administration, supervision, funding acquisition, methodology, conceptualization, writing – original draft and review & editing.

## Data availability

Methods and supplemental tables and figures are available as part of the ESI. Raw data files are available in a dataset in the Harvard Dataverse.

## Acknowledgments

This work was funded by the National Institute of Health grants R01GM132564 (E.P.B.) and P01 GM077596 (W.v.d.D.). G.E.K. was a Merck Fellow of the Damon Runyon Cancer Research Foundation, funded by DRG2369-19. E.P.B. and W.v.d.D. are HHMI Investigators. Metal analysis was performed at the Northwestern University Quantitative Bio-element Imaging Center generously supported by NASA Ames Research Center NNA06CB93G. We thank the Laukien-Purcell Instrumentation Center at Harvard University provided access to NMR instrumentation. Access to the Bruker EPR was kindly provided by the Department of Chemistry Instrumentation Facility (DCIF) at the Massachusetts Institute of Technology. This article is subject to HHMI’s Open Access to Publications policy. HHMI lab heads have previously granted a nonexclusive CC BY 4.0 license to the public and a sublicensable license to HHMI in their research articles. Pursuant to those licenses, the author-accepted manuscript of this article can be made freely available under a CC BY 4.0 license immediately upon publication.

