## Supplemental Information for "A new heme enzyme family forms hydrazine groups in diverse biosynthetic pathways"

**Electronic Supplementary Information for "A new heme enzyme family forms hydrazine groups in diverse biosynthetic pathways"**

Grace E. Kenney[1,‡], Kwo-Kwang A. Wang[2,‡], Tai L. Ng[1,‡], Wilfred A. van der Donk[2,3,4], Emily P. Balskus[1,5*]
[1] Department of Chemistry and Chemical Biology, Harvard University, Cambridge 02138 MA, USA
[2] Department of Chemistry, University of Illinois at Urbana-Champaign, Urbana 61801 IL, USA
[3] Carl R. Woese Institute for Genomic Biology, University of Illinois at Urbana-Champaign, Urbana 61801 IL, USA
[4] Howard Hughes Medical Institute, University of Illinois at Urbana-Champaign, Urbana 61801 IL, USA
[5] Howard Hughes Medical Institute, Harvard University, Cambridge 02138 MA, USA

### Table of Contents

**Supplementary Files**
All supplemental files will be made fully public as a Harvard Dataverse dataset upon publication.

**File SF01:** KinJ_gene_seqs.fa (.fasta file)
**File SF02:** KinJ_gene_metadata_full.xlsx (Excel file)
**File SF03:** KinJ_neighbor_seqs.fa (.fasta file)
**File SF04:** KinJ_neighbor_metadata_full.xlsx (Excel file)
**File SF05:** KinJ_ssn-gnn.cys (Cytoscape file)
**File SF06:** Fdx-ssn.cys (Cytoscape file)
**File SF07:** Fdx_ssn_metadata.xslx (Excel file)
**File SF08:** KinJ-related_constructs.gb (multi-Genbank file)
**Files SF09a-c:** KinJ SDS-PAGE (Coomassie), Western (anti-$His_6$-AP), and Coomassie vs. TMBZ stain (.tiff for gels and .jpg for Western photographs)
**Files SF10a-b:** KinI SDS-PAGE (Coomassie) and Western (anti-$His_6$-HRP) (.tiff for gels and .jpg for Western photographs)
**Files SF11a-b:** KinIJ SDS-PAGE (Coomassie) and Western (anti-$His_6$-AP) (.tiff for gels and .jpg for Western photographs)
**Files SF12a-b:** FzmP SDS-PAGE (Coomassie) and Western (anti-$His_6$-HRP) (.tiff for gels and .jpg for Western photographs)

**Supplementary Methods**
Pg. 3 Bioinformatics
Pg. 5 Cloning and mutagenesis
Pg. 6 Protein expression and purification
Pg. 8 Electronic absorption spectroscopy
Pg. 9 Evaluation of metal loading
Pg. 9 Heme extraction
Pg. 10 Electron paramagnetic resonance spectroscopy
Pg. 10 Synthesis of L-hydrazinosuccinate standard
Pg. 12 Kin(I)J activity assays

**Supplementary Figures**
Pg. 14 **Figure S1:** Enzymatic origins for N–N bonds.
Pg. 15 **Figure S2:** KinJ is a heme protein.

Pg. 16 **Figure S3:** KinI is a 2[4Fe-4S] ferredoxin.
Pg. 17 **Figure S4:** KinIJ co-expression confirms heterodimer formation.
Pg. 18 **Figure S5:** L-Hzs synthesis.
Pg. 19 **Figure S6:** Mass spectrometry for L-Asp and L-Hzs standards.
Pg. 20 **Figure S7:** Further exploring KinJ activity.
Pg. 21 **Figure S8:** KinJ versus KinJ activity
Pg. 22 **Figure S9:** $^{15}N$ fragmentation confirms location of the $NO_2^-$-derived nitrogen in L-Hzs.
Pg. 23 **Figure S10:** Prioritizing KinJ variants
Pg. 24 **Figure S11:** KinJ variants point towards a possible axial ligand and active site.
Pg. 26 **Figure S12:** Models for the KinJ fold are consistent across methods and oligomeric states.
Pg. 27 **Figure S13:** The KinJ active site is in a C-terminal domain with structural homology to the GAF domain superfamily.
Pg. 29 **Figure S14:** The identities of proposed 4Fe-4S ferredoxin partners differ, but their binding sites remain consistent.
Pg. 31 **Figure S15:** Structural models of binding of ferredoxin partners to KinJ homologs

**Supplementary Tables**

Pg. 33 **Table S1:** Organisms
Pg. 34 **Table S2:** Primers
Pg. 35 **Table S3:** Constructs
Pg. 36 **Table S4:** KinJ DALI & Foldseek results

**Supplementary References**

Pg. 38

**Supplementary Methods**

Unless otherwise specified, chemicals were purchased from MilliporeSigma or Fisher Scientific and were ACS or Molecular Biology grade. $^{15}N$-labeled chemicals were purchased from Cambridge Isotopes. Strains are listed in **Supplementary Table S1.**

*Bioinformatics*

KinJ sequences do not belong to any pre-existing protein families, and *kinJ* homologs are generally annotated solely as hypothetical proteins. The *S. murayamaensis* sequence was thus used as the seed for an initial protein BLAST search against the IMG/M genome data base as well as the NCBI Refseq database.[1] The most divergent result was then used to initiate a second BLAST round. This process was repeated until no additional KinJ sequences were detected, and the results were pooled. The resulting 1042 amino acid sequences and metadata of the genes encoding these homologs were downloaded on 2023.08.05 and are available as supplementary files, with additional information added to the metadata as described below. Prior to further analysis, KinJ sequences were aligned using MAFFT (L-INS-I mode)[2] and truncated KinJ sequences with under 450 amino acids were removed from sequence-based analyses.

The prettyClusters R package was used to analyze genome neighborhood content, to identify subgroups of neighboring protein families, to identify subgroups of similar genome clusters, and to visualize genome neighborhoods.[3] Genes within a 10-gene range up- and down-stream from *kinJ* homologs were considered for genome neighborhood analysis. Sequences derived via NCBI were reannotated using InterProScan[4,5] prior to their addition to the IMG-derived dataset, and ferredoxins in *Corynebacterium* BGCs were manually annotated. Analyses were run with the full KinJ dataset and with representative nodes from the KinJ SSN only, and were run with and without removal of truncated BGCs located near a contig end.

The trimmed dataset – with truncated contigs removed – consisted of 869 KinJ sequences and their genomic neighborhoods. A sequence similarity network (SSN) was constructed for KinJ sequences using the EFI-EST web tool,[6,7] employing an e-value cutoff of 1E-65 and a representative node cutoff of either 95 or 100% identity. This set of 234 representative nodes – corresponding to specific KinJ sequences – and their genomic neighborhoods were used for the sequence similarity and genome similarity network analysis shown in this paper. The prettyClusters R package was used to provisionally assign types of genome neighborhoods and construct a genome neighborhood network,[3] and Cytoscape was used to visualize both sequence and genome neighborhood networks.[8] Provisional groups of hypothetical proteins were assigned, including a subset of unannotated KinI-like ferredoxins,

In the genomes of 149 species lacking a ferredoxin encoded within 10 genes of a KinJ homolog, sequences for proteins encoded elsewhere that contained one or more 3Fe-4S or 4Fe-4S clusters were identified via InterPro annotations; sequences over 150 aa were filtered out. An SSN was constructed for the remaining sequences as described above, but using an e-value cutoff of 1E-41 and no representative nodes; two clusters (one widespread) with homologs in some KinJ-encoding BGCs were detected. Representative members of KinJ-associated ferredoxins were aligned with selected characterized ferredoxins using MAFFT[2] under E-INS-I mode.

MEGA version 12 was used to identify the best amino acid substitution models (WAG with a gamma distribution and invariant sites).[9] The gamma distribution was modeled across 5 categories (+G, parameter = 1.9839) with 4.09% of sites determined to be evolutionarily invariant in this dataset (+I). Phylogeny was inferred using an adaptive bootstrap approach as the phylogenetic test for the maximum likelihood analysis of the dataset. Initial neighbor-joining trees were constructed using a matrix of pairwise distances computed using the p-distance. Initial maximum parsimony trees were constructed using the tree with the shortest length among 10 MP searches performed with a randomly generated starting tree. The tree with the lowest log-likelihood score between the initial NJ and MP trees was used for further maximum likelihood analyses. The adaptive bootstrap test used with a threshold of 5.0, and 104 replicates were ultimately run to generate the final bootstrapped consensus tree. For alignment visualization, MAFFT-aligned single representatives of ferredoxin classes were compared to KinJ-associated ferredoxin sequences.

Protein structure prediction was carried out using AlphaFold2 Multimer (for apo structures) and AlphaFold3 (for heme-bound structures).[10,11] Some predictions used the ColabFold implementation of AlphaFold2 Multimer.[12] KVfinder and Caver were used to identify cavities,[13,14] and PyMOL (Version 3.0, Schrödinger LLC) was used to align protein structures and to generate charge surface views via the Adaptive Poisson-Boltzmann Solver plugin. The AlphaFold-predicted structures were submitted to the DALI and Foldseek protein structure comparison servers to identify remote homology.[15,16]

*Cloning and mutagenesis*

Actinomycetes – including *Kitasatospora purpeofusca, Streptomyces murayamaensis, Streptomyces* sp. S-1521, *Streptomyces* sp. S-149, and *Streptomyces* sp. XY332 were grown in 50 mL ISP-2 or R2B medium at 30 °C, shaking at 180 rpm for 1 week. 1 mL of cells (pelleted at 20k x g and 4 °C for 15 min) or a similar amount of flocculant cells were used for genomic DNA isolation via a MasterPure GramPositive kit (Epicentre).

Primers were synthesized by MilliporeSigma. For amplification of nucleic acids from plasmid constructs or from purified PCR products, Q5 polymerase (NEB) was used. For amplification of nucleic acids from gDNA, PrimestarGXL (Takara) was used. Restriction enzymes and T4 DNA Ligase were purchased from NEB and used as directed; where specified, InFusion kits (Takara) were used for construct assembly instead. Plasmids were amplified in the Top10 (Invitrogen) or *E. cloni* 10G (Lucigen) *E. coli* strains. DNA cleanup, gel extraction, and miniprep isolation of plasmids was performed using kits produced by Zymo. Nucleic acids were quantified via NanoDrop 1000 (Thermo). Sequencing for all constructs in pET-28a and pET-15b was performed using the T7F and T7R primers by Azenta Life Sciences. Sequencing for all constructs in Duet vectors was performed using the ACYCDuet-UP1,

DuetDOWN-1, DuetUP2, and T7R primers by Azenta Life sciences; in pET vectors, sequencing used the T7F and T7R primers.

Constructs for heterologous expression of *Streptomyces murayamaensis* KinJ, *S.* sp. S-149 FzmP, *S.* sp. XY332 FzmP, and *S.* sp. XY332 FzmQ were designed using primers listed in **Supplementary Table 2**, and cloned into pET-28a (KinJ) or pET-15b (FzmP, FzmQ) (Novagen) via restriction digestion and ligation. These constructs contained an N-terminal $His_6$ tag followed by a thrombin cleavage site. Several variants for *Smu*-KinJ (Y346F, C468A, M498A, H504A) were generated via a QuikChange Lightning kit (Agilent) as directed, using primers listed in **Supplementary Table 2**. Additional variants (Y223F, Y304F, K356A, M436A, H537A, H541A, and H541C) were synthesized and cloned by Genscript.

Co-expression constructs for pairs of KinJ homologs and their redox partners were established in pCDFDuet-1 (Novagen). Co-expression constructs for *Smu*-KinI and *Smu*-KinJ homologs from *S. murayamaensis, S.* sp. S-149, and *S.* sp. XY332 were designed using primers listed in **Supplementary Table 2**, and cloned into pCDFDuet-1, with N- or C-terminal $His_6$ or Strep tags (and enterokinase cleavage sites) as specified; other constructs in **Supplementary Table 3** used codon-optimized sequences were synthesized and cloned into pCDFDuet-1 by Genscript. Unless otherwise specified, in these constructs, KinJ homologs contained an N-terminal $His_6$ affinity tag, followed by an enterokinase cleavage site and then the authentic N terminus of the protein. While FzmP-containing constructs from two species were tested, results shown in this manuscript are from the *S.* sp. S-149 homolog unless otherwise specified. GenBank-formatted sequences for all constructs are listed in **Supplementary Table 3** and are available as a supplemental multiGenBank file.

*Protein expression and purification*

All enzymes were expressed using minor variations on the same protocol. The plasmids were used to transform chemically competent LOBSTR (DE3) (Kerafast) [17]. 5 mL cultures in LB with the appropriate antibiotics (predominantly kanamycin at 50 μg/mL or spectinomycin at 70 µg/mL) were grown overnight at 37 °C and 180 rpm, and in the morning, 1 mL of the overnight culture was used to inoculate a 100 mL starter culture in LB. This starter culture was used to inoculate 4-6 L of autoinduction media, which was amended with 250 µM ferrous ammonium sulfate and 250 µM δ-aminolevulinic acid during the production of KinJ homologs and variants, and 500 µM ferrous ammonium sulfate and 250 µM cysteine during the production of KinI homologs. Large-scale cultures were incubated at 37 °C and 200 rpm for 4 h, followed by a reduction in temperature to 18 °C. Growth continued overnight, and cells were harvested after ~18 h by centrifugation at 8000 x g and 4 °C for 10 min. Cell pellets were frozen in liquid nitrogen and stored at –80 °C until purification.

All aerobically purified KinJ homologs were purified following the same protocol. Buffers were prepared, filtered through 0.2 µm membranes, and chilled at 4 °C. The cell pellets were resuspended in 50 mL of lysis buffer (50 mM MOPS pH 7.25, 250 mM NaCl, 10% glycerol, 10 mM imidazole) supplemented with ~ 1 mg/mL DNase I (MilliporeSigma Sigma) and half of a Sigma EDTA-free protease inhibitor tablet (MilliporeSigma). Cells were lysed on ice via sonication at 25% power using a Sonifier (Branson) for 7.5 min, with 1 s pulses every 4 s. The cell lysate was clarified via centrifugation at 20 k x g and 4 °C for 45 min. The resulting supernatant was applied at 4 °C to a Ni-loaded 5 mL HisTrap column (Cytiva) pre-equilibrated with lysis buffer, using a Bio-Rad DuoFlow II FPLC system (Bio-Rad). The flow-through was discarded and the resin was washed with 10 column volumes of lysis buffer. The $His_6$-tagged enzymes were eluted over a gradient of 10-500 mM imidazole.

Fractions were analyzed using sodium dodecyl sulfate-polyacrylamide gel electrophoresis (SDS-PAGE) using 16% or 10-20% precast Novex gels (Thermo), and – when required – confirmed via Western blots, using PVDF membrane, anti-$His_6$-HRP conjugate antibodies (MilliporeSigma) and a colorimetric CN-DAB detection kit (Pierce). Fractions containing the protein of interest were pooled, concentrated via an Amicon spin concentrator with a 30 kDa MWCO membrane, and loaded onto a pre-equilibrated and calibrated 120 mL HiLoad Superdex 200 16/600 (homologs of KinJ and KinIJ co-expression constructs) or HiLoad

Superdex 75 16/600 (KinI, FzmQ) size exclusion column (Cytiva) and eluted over 1.3 column volumes. Fractions were again analyzed using SDS-PAGE and Western blot, and fractions were concentrated and quantified via Bradford assay (Bio-Rad) or via predicted extinction coefficient and measurement of absorbance at 280 nm. Where appropriate, samples were assayed for cofactor loading as described subsequently, and then flash frozen in liquid nitrogen and stored at –80 °C. If used for anaerobic experiments, the protein was deoxygenated under alternating vacuum and inert gas on a Schlenk line, sealed, and brought into the anaerobic chamber for further use.

All anaerobic KinI, KinJ, and KinIJ constructs were purified via a similar protocol. However, for anaerobic protein purification, all buffers were deoxygenated and brought into an anaerobic chamber (Coy) maintained at 4 °C; buffers and plasticware were introduced to the chamber a day before use. Cell pellets were thawed and lysed in the anaerobic chamber, and the sonification pulse sequence was decreased to 1 s on and 6 s off. Cell lysate was removed from the anaerobic chamber in sealed Nalgene Oak Ridge 42 mL round-bottom tubes with rubber O-ring-lined caps, clarified as described previously, and re-introduced into the anaerobic chamber. Lysate was loaded on an anaerobic Bio-Rad DuoFlow FPLC as described above, and fractions were concentrated in Amicon spin concentrators with a 3 kDa MWCO membrane (samples with KinI) or a 30 kDa MWCO membrane (samples with only KinJ homologs) in a refrigerated centrifuge in an anaerobic chamber. Size exclusion chromatography was also performed anaerobically but otherwise as described above, and concentrated samples were sealed, flash frozen in liquid nitrogen, and stored in a liquid nitrogen storage dewar until use.

*Electronic absorption spectroscopy*

Aerobic optical spectra were collected using samples at 25 μM enzyme concentration (unless otherwise noted) in an ultra-low volume quartz cuvette (Hellma.) Samples were collected at room temperature or at 10 °C in an Agilent 8454 spectrophotometer (Agilent). Anaerobic spectra were collected similarly except that the cuvette was loaded in an anaerobic chamber. In the absence of an anaerobic UV-vis, the cuvettes were sealed with a rubber septum, and when substrates or other reagents were introduced, they were added using a gas-tight Hamilton syringe. For protein concentration determination via absorbance at A280, the extinction coefficients predicted via ProtParam (Expasy) were used. For protein concentration determination via Bradford assay, that Bio-Rad reagent was used. In both cases, multiple dilutions were measured to reach the linear range of detection. For evaluation of cofactors, samples were diluted to target an absorption of 0.1-1 AU at the wavelength of interest (generally 10-20 µM for KinJ homologs at the Soret peak.) The pyridine hemochromagen assay was performed as described previously.[18]

*Evaluation of metal loading*

For metal analysis, all samples were digested in 5% trace metal grade nitric acid and 0.75% trace metal grade $H_2O_2$ in metal-free tubes (VWR) and were heated to 60 °C during digestion. Sample dilutions targeted the 10-200 ppb range, and if necessary, samples were pre-diluted so that the added protein volume did not exceed 1:100 of the final sample volume. All samples – including a buffer-only blank - were run in triplicate. Standard curves were generated via a dilution series of a multi-element standard containing 10 µg/mL (10 ppm) each As, Ca, Cd, Co, Cr3, Cu, Fe, K, Mg, Mn, Mo, Ni, Se, V, and Zn in 2% $HNO_3$ (v/v) with trace HF, with density at 1.011g/mL (Inorganic Ventures). The metal content of the samples was measured using a Thermo iCAP ICP-MS instrument in the Quantitative Bio-element Imaging Center (QBIC) core facility at Northwestern University. The standard curve was used to calculate metal concentrations and stoichiometry.

*Heme extraction*

Heme extraction was performed on 150 µL of 100 µM enzyme. To these samples, 3 µL of concentrated HCl was added to precipitate the protein. Samples were vortexed for 1 min and 2 volumes (300 µL) ethyl acetate was added. Samples were again vortexed for 1 min and then centrifuged at 13k x g rpm at 4 °C. The organic fraction was removed via pipet and dried *in vacuo.* Samples were resuspended in 100% MeOH and then analyzed via UV-visible light spectroscopy or LC-MS. During LC-MS, samples were diluted 1:10 in water, and 10 µL samples were injected onto a Dikma BioBond C4 column (5 µ, 4.6 i.d. x 50 mm). Samples were eluted using a gradient from 5% to 95% solvent B (0.1% formic acid in acetonitrile) in solvent A (0.1% formic acid in water) over 15 min at .5 mL / min on an Agilent 1200 series LC system. Mass spectrometry was performed using an Agilent Q-TOF 6530 with an ESI source. The following parameters were used for the Q-TOF: drying gas 8 L/min, gas temperature 325 °C, fragmentor voltage 125 V, nebulizer pressure 35 psi, capillary voltage 3500 V, nozzle voltage 1000 V. MS1 spectra were acquired at 1 spectrum/s with a minimum m/z of 100 Da. Mass spectra and extracted ion chromatograms were exported using MassHunter (Agilent). Extracted ion chromatograms by default used a 10 ppm cutoff.

*Electron paramagnetic resonance spectroscopy*

EPR samples were prepared in 50 mM MOPS pH 7.25, 250 mM NaCl, and 10% glycerol. 250 µL samples were loaded into Wilmad quartz X-band EPR tubes (Wilmad) and samples were flash-frozen in liquid nitrogen (aerobically or anaerobically), where they were stored until analysis. Continuous wave (CW) X-band spectra were collected on a Bruker EMX-Plus spectrometer, cooled to 10 K via a Cturker/ColdEdge 4K crytogen-free cryostat. Data were analyzed using EasySpin.[19]

*Synthesis of* L*-hydrazinosuccinate standard*

Synthesis of L-hydrazinosuccinate utilized an oxaziridine reagent to generate the hydrazine from the corresponding amine (**Fig. S9A**), following a route established by Kang et al.[20] Synthesis of 2-*tert*-butyl 3,3-diethyl 1,2-oxaziridine-2,3,3-tricarboxylate was carried out as described in that publication.

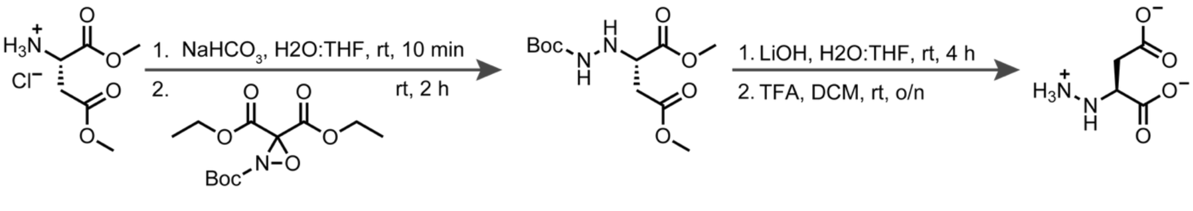

Commercially available L-aspartate dimethyl ester hydrochloride (Chem-Impex) provided the starting point for L-hydrazinosuccinate synthesis. L-aspartate dimethyl ester hydrochloride (100 mg, 0.506 mmol, 1 equiv) was resuspended in 5 mL of THF and 5 mL of saturated aqueous sodium bicarbonate. The resulting mixture was stirred for 10 min at room temperature. 1.1 equivalents of the oxaziridine reagent (161 mg, 0.557 mmol)) were added, and the reaction mixture was stirred for 2 h at room temperature. After completion of the reaction, 10 µL of ethylenediamine (.3 equiv) was added, followed by 10 mL of ethyl acetate and 20 mL of 1M HCl. To remove the amine starting material, the organic layer was washed with 3 x 10 mL of 1 M HCl and 10 mL of brine. The organic layer was dried over anhydrous sodium sulfate and concentrated *in vacuo*, generating the Boc-protected hydrazinodimethyl ester which was used without further characterization.

To saponify the methyl ester groups, the crude dimethyl ester product was dissolved in 5 mL of THF and 5 mL of 1 M LiOH (aq), and the reaction mixture was stirred at room temperature. After 5 h, the mixture was diluted with 10 mL of water and washed with 2 x 10 mL ether to remove remaining dimethyl ester. The pH of the aqueous layer was adjusted to 1 with concentrated HCl, and the Boc-protected hydrazino acid was extracted into 2 x 10 mL of ethyl acetate. The organic layers were combined, washed with 1 x 10 mL of brine, dried over sodium sulfate, and concentrated *in vacuo*. The crude product was resuspended directly in 25% TFA in dichloromethane and was allowed to stir at room temperature overnight. TFA and DCM were removed via rotovap, and the resulting oil was triturated with ether to afford crude L-hydrazinosuccinate.

The crude L-hydrazinosuccinate was dissolved in water, and the mixture was purified via semi-preparative reverse-phase HPLC on a Dionex Ultimate 3000 (Thermo Scientific). The LC column was a Kromasil 100 C18 column (250 x 10 mm, 5 µ). The LC solvents were 0.1% formic acid in water (A) and 0.1% formic acid in acetonitrile (B). After application to the column, the LC program began with a shallow gradient of 5-12 % B over 10 min, followed by a linear gradient from 12-30% B for 2.5 min, a linear gradient from 30% to 5% B in 12.5 min, and an isocratic hold at 5% B for 3 min, all at a constant flow rate of 3 mL/min. Elution of the compound was monitored at 200 nm. The collected fractions were lyophilized, producing a white solid (~55 mg, ~75 % yield) which was resuspended in $D_2O$ for initial analysis. After NMR characterization, the solution was lyophilized and resuspended in 250 µL water, producing a 10 mM stock solution of L-hydrazinosuccinate.

NMR spectra were obtained on a Varian Mercury 400 (400 MHz). The chemical shifts (δ) are reported in parts per million (ppm) downfield from tetramethylsilane, and the solvent resonance is used as an internal standard ($D_2O$ = 4.79 ppm). $^1$H NMR for L-hydrazinosuccinate (400 MHz, 100% $D_2O$): δ (ppm) ~ 3.8 (t, 1H, $H_{12}$), ~2.8 (dd, 1H, $H_{11}$), ~2.6 (dd, 1H, $H_{10}$) (**Fig. S9B**). $^1$H NMR for L-hydrazinosuccinate (400 MHz, 10% $D_2O$): δ (ppm) ~4.2 (t, 1H, $H_{12}$), ~3.8 (m, 1H, $H_{13}$), ~2.9 (dd, 1H, $H_{11}$), ~2.8 (m, 2H, $H_{14}$ & $H_{15}$), ~2.7 (dd, 1H, $H_{10}$) (**Fig. S9C**). HRMS for L-hydrazinosuccinate $[M-H]^-$ ($C_4H_7N_2O_4^-$): calc. 147.0411 *m/z*, obs. 147.0404.

*Kin(I)J activity assays*

Assays to measure the activity of KinIJ homologs were carried out in an anaerobic chamber at 4 °C. All stocks were freshly prepared in deoxygenated reaction buffer. The standard reaction buffer consisted of 50 mM MOPS pH 7.25, 100 mM $NH_4Cl$, and 10% glycerol. Reaction mixtures contained a final 100 µM KinJ or KinIJ, 250 µM amino acid substrate (most commonly L-Asp, prepared as a 2.5 mM stock), 500 µM nitrogen substrate (most commonly $NaNO_2$, prepared as a 10 mM stock), 5 mM sodium dithionite, and 1.5 mM electron mediator (methyl viologen or benzyl viologen unless otherwise specified.) Reactions were carried out over 12 h unless otherwise specified, and either quenched with oxygen for immediate processing or frozen in liquid nitrogen.

For all KinIJ activity assays, proteins were removed via centrifugation using a 3 kDa MWCO spin concentrator (Amicon). The eluate was diluted 1:10 in water, and 20 µL samples were maintained in a chilled autosampler at 8 °C until they were injected onto a HydroBondAQ column (Mac-Mod: 5 µ, 250 x 4.6 mm i.d.) Samples were eluted using a gradient from 100%

solvent A (0.1% formic acid in water) to 35% solvent B (0.1% formic acid in acetonitrile) over 15 min at 0.5 mL/min on an Agilent 1200 series LC system. Mass spectrometry was performed using an Agilent Q-TOF 6530 equipped with an ESI or Dual AJS ESI source. The following parameters were used for the Q-TOF when using an ESI source: drying gas 8 L/min, gas temperature 300 °C, fragmentor voltage 125 V, nebulizer pressure 35 psi, capillary voltage 3500 V, nozzle voltage 1000 V. When using a Dual AJS source, drying gas flow was at 11 L/min, and the following additional parameter were used: sheath gas flow 11 L/min, sheath gas temperature 325 °C. With both instrument configurations, data collection used negative ion mode and MS1 spectra were collected at a rate of 1/s with a mass range of 70-1700 m/z. For some substrates, a second run in positive ion mode was added (with expected MS1 values adjusted accordingly). Targeted MS2 fragmentation was performed with a minimum MS and MS/MS *m/z* of 50 Da, acquisition at 1 spectrum/s and a medium isolation width, with a default collision energy of 10 V. Mass spectra and extracted ion chromatograms were exported using MassHunter (Agilent). Extracted ion chromatograms by default used a 10 ppm cutoff.

### Supplementary Figures

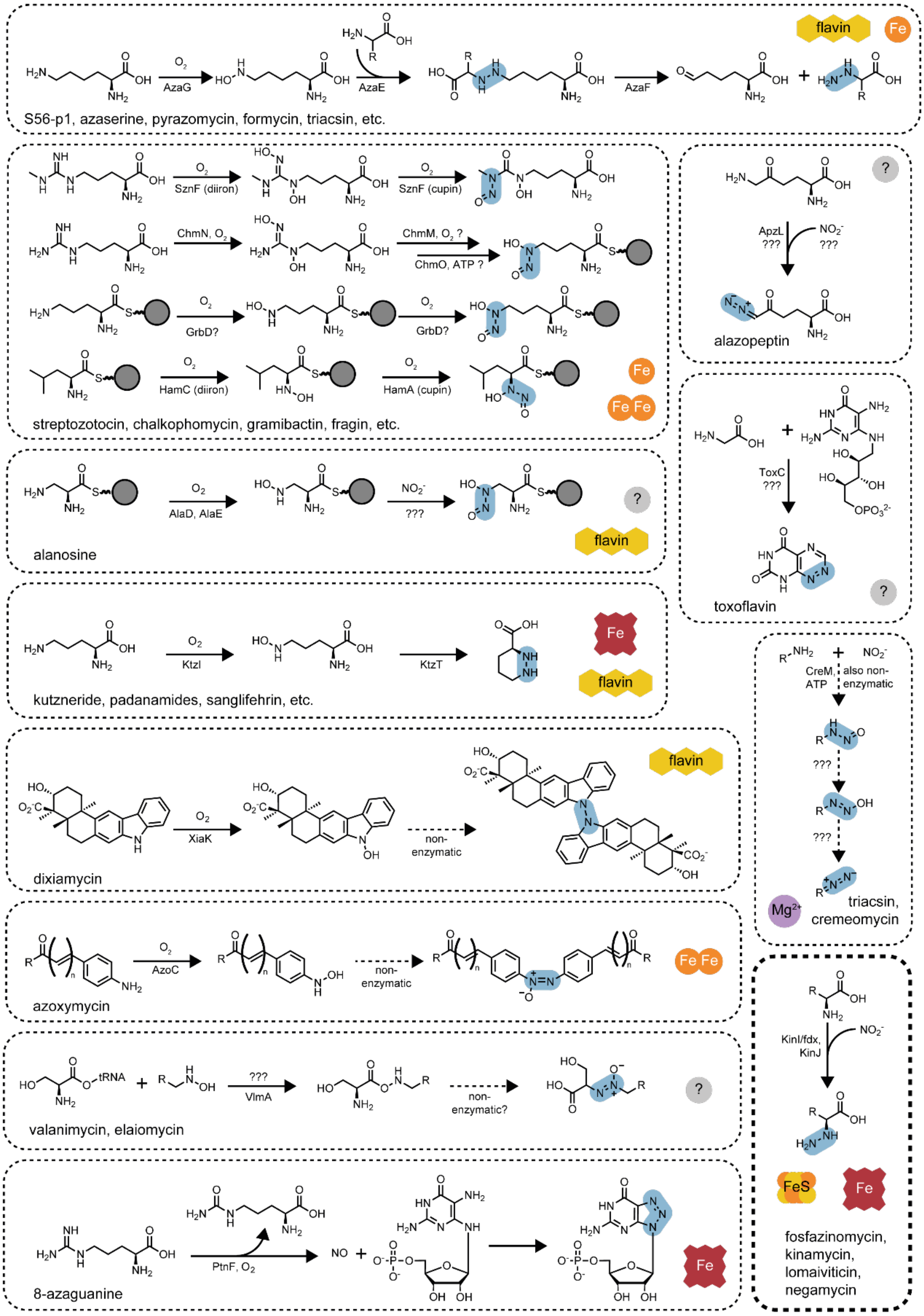


**Figure S1. Enzymatic origins of N–N bond formation in natural product biosynthesis.** With the identification of KinIJ, at least a dozen distinct enzymatic systems for N–N bond biosynthesis in natural products have been explored. One particularly common biosynthetic strategy relies on the oxidation of one amine substrate to form a hydroxylamine, activating it for N–N bond formation with an amine co-substrate.

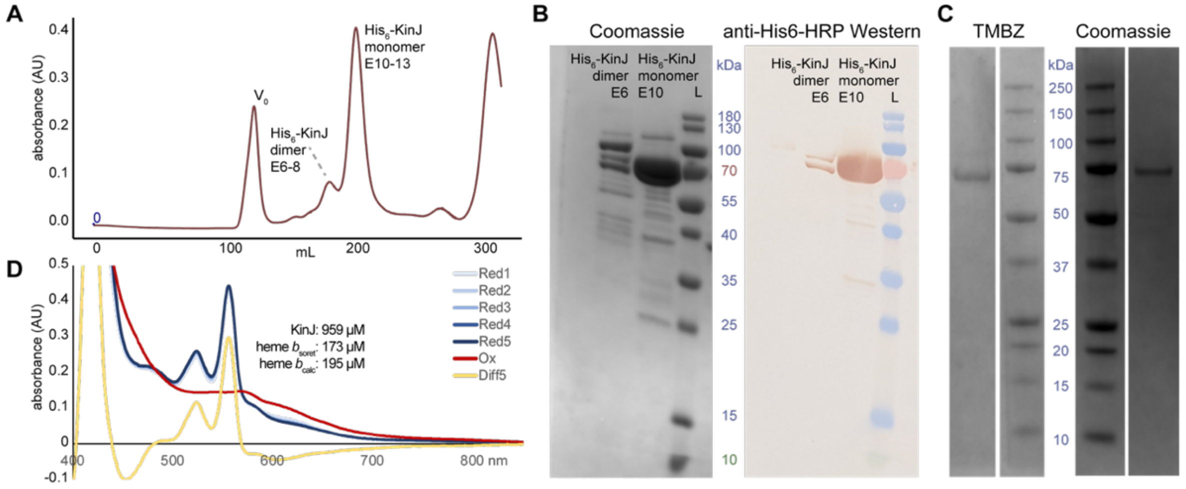


**Figure S2. KinJ is a heme enzyme**. **A**) Superdex 200 26/600 size exclusion chromatogram for His$_6$-Thr-KinJ. A small peak consistent with a homodimer and a predominant monomer form are observed. **B)** SDS-PAGE (16% Novex Tris-glycine) and anti-His$_6$-HRP Western blot (visualized with CN/DAB) for His$_6$-Thr-KinJ. **C)** In-gel TMBZ assay for His$_6$-Thr-KinJ demonstrates peroxidase activity at a mass consistent with KinJ, supporting the identification of KinJ as a heme enzyme. The in-gel assay was performed at 5x higher concentration; standard loading was used for the Coomassie-stained comparison. **D**) Heme loading estimates via absorption at the oxidized Soret peak using the extinction coefficient of myoglobin heme *b* could be benchmarked using the pyridine hemochromogen assay.[21] In this example during early attempts to quantify heme loading *in vivo*, observed heme loading was ~20%.

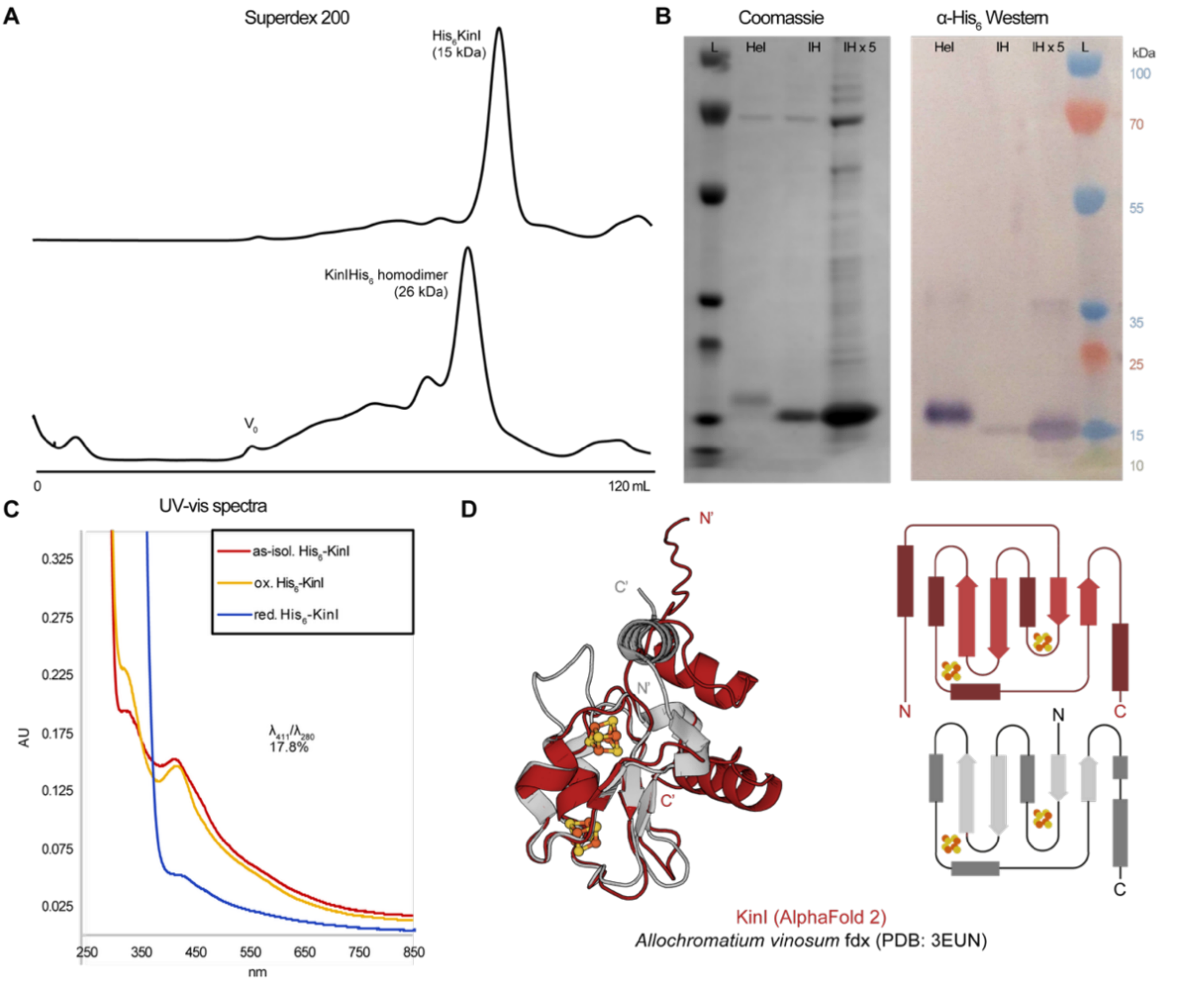


**Figure S3. Purification of KinI. A**) Superdex 200 size exclusion chromatogram for $His_6$-Ek-KinI and KinI-$His_5$. Note that later versions of this construct added an additional C-terminal His. **B**) Coomassie stained gel and anti-$His_6$ Western blot for heterologously expressed KinI. **C**) UV-vis spectra of KinI are consistent with a 2[4Fe-4S]$^{2+/+}$ ferredoxin. **D**) AlphaFold2 model of KinI (red) aligned against the structure of a 2[4Fe-4S]$^{2+/+}$ ferredoxin from *Allochromatium vinosum* (PDB: 3EUN, grey), the prototypical "Alvin" ferredoxin[22] with an RMSD of 1.508 Å. The core ferredoxin fold is highly conserved. However, compared to Alvin ferredoxins, KinI has an N-terminal extension with some alpha helical regions. In the AlphaFold2 model, this extension displaces the C-terminal helix common to both ferredoxins. Additionally, the C-terminal 4Fe-4S site (cluster I) contains a standard $Cx_2Cx_2Cx_3C$ motif, rather than the lengthier $Cx_2Cx_8Cx_3C$ motif found in Alvin ferredoxins.[23]

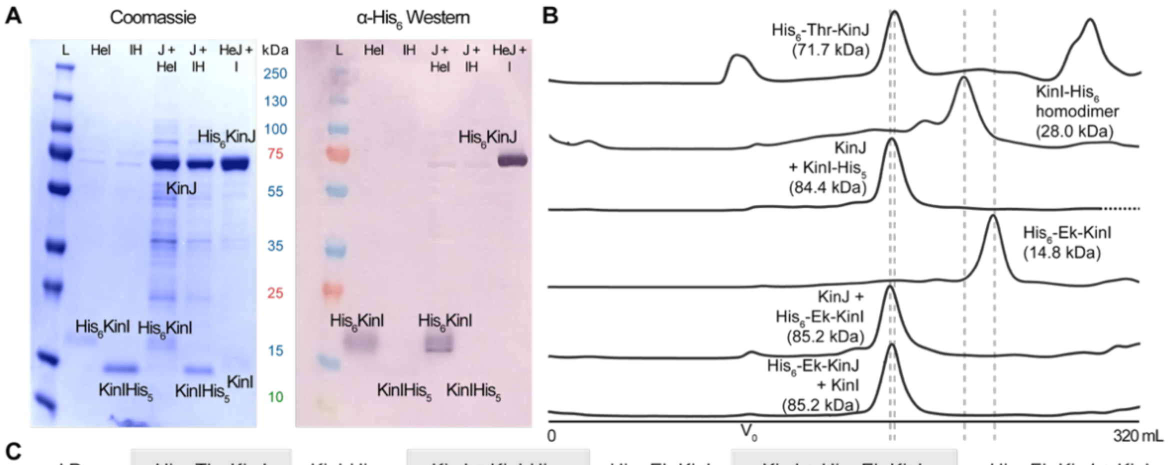


C

| kDa | $His_6$-Thr-KinJ | KinI-$His_6$ | KinJ + KinI-$His_6$ | $His_6$-Ek-KinI | KinJ + $His_6$-Ek-KinI | $His_6$-Ek-KinJ + KinI |
|---|---|---|---|---|---|---|
| $pred_{mono}$ | 71.7 | 14.0 | n/a | 14.8 | n/a | n/a |
| $pred_{dimer}$ | 143.4 | 28.0 | 84.4 | 29.6 | 85.2 | 85.2 |
| obs | 83.7 | 19.8 | 91.1 | 9.5 | 89.6 | 86.4 |

**Figure S4. Co-expressed KinI and KinJ form a heterodimer. A**) SDS-PAGE and anti-$His_6$ Western blot for singly and co-expressed KinI and KinJ (isolated anaerobically). Whether the $His_6$ tag is located on KinI or KinJ, each protein can pull the other down. **B**) Size exclusion chromatography for co-expressed KinIJ supports heterodimer formation. Depicted KinI, KinJ, and KinIJ complexes were analyzed on a Superdex 200 26/600 column, and all traces are normalized by maximum absorbance at 280 nm. **C**) After calibration against gel filtration standards (Bio-Rad), calculated masses were compared to predicted masses to ascertain the oligomeric state.

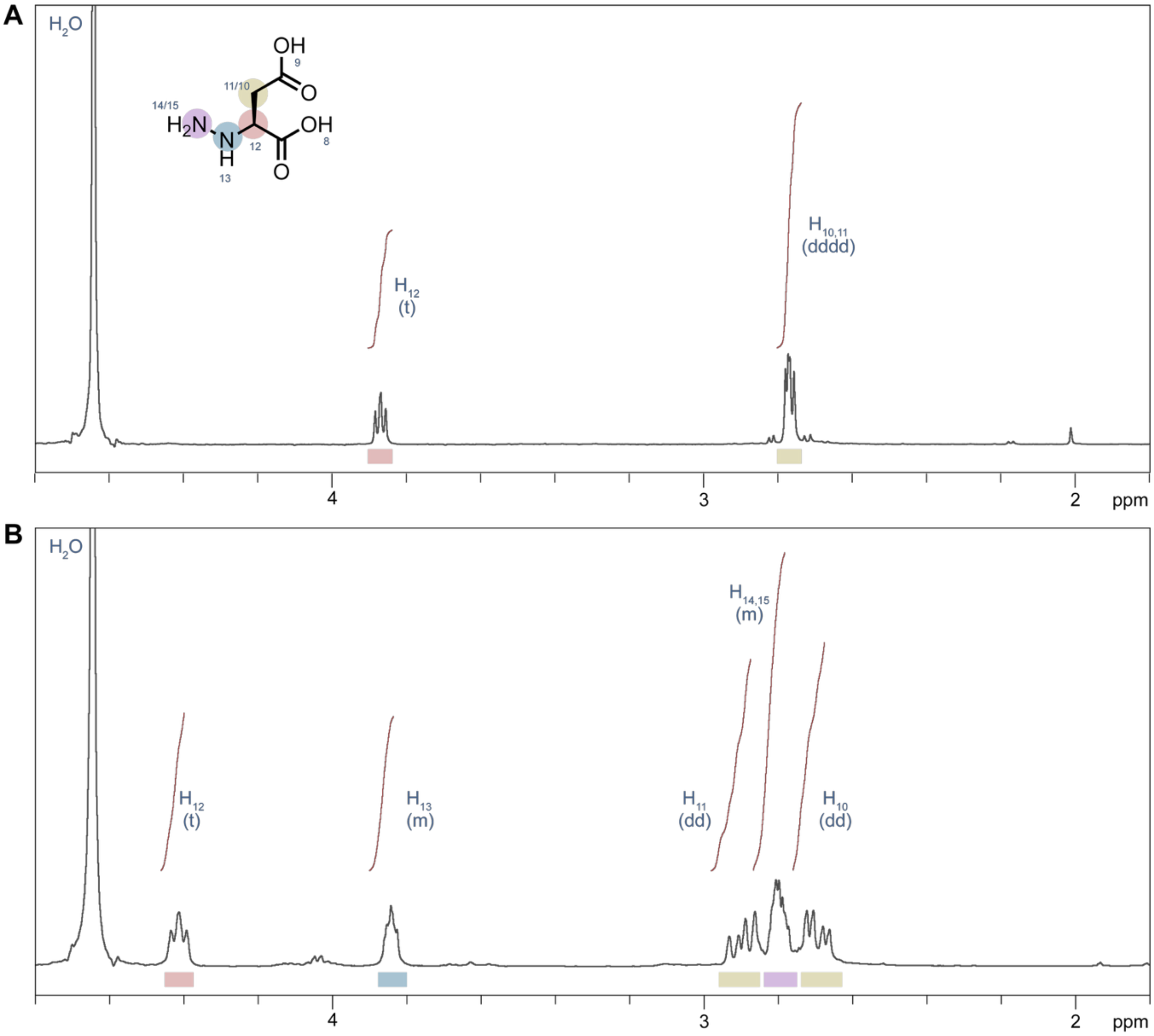


**Figure S5. [1]H NMR for an L-hydrazinosuccinate standard. A)** [1]H NMR of L-hydrazinosuccinate in $D_2O$. **B)** [1]H NMR of L-hydrazinosuccinate in 10% $D_2O$.

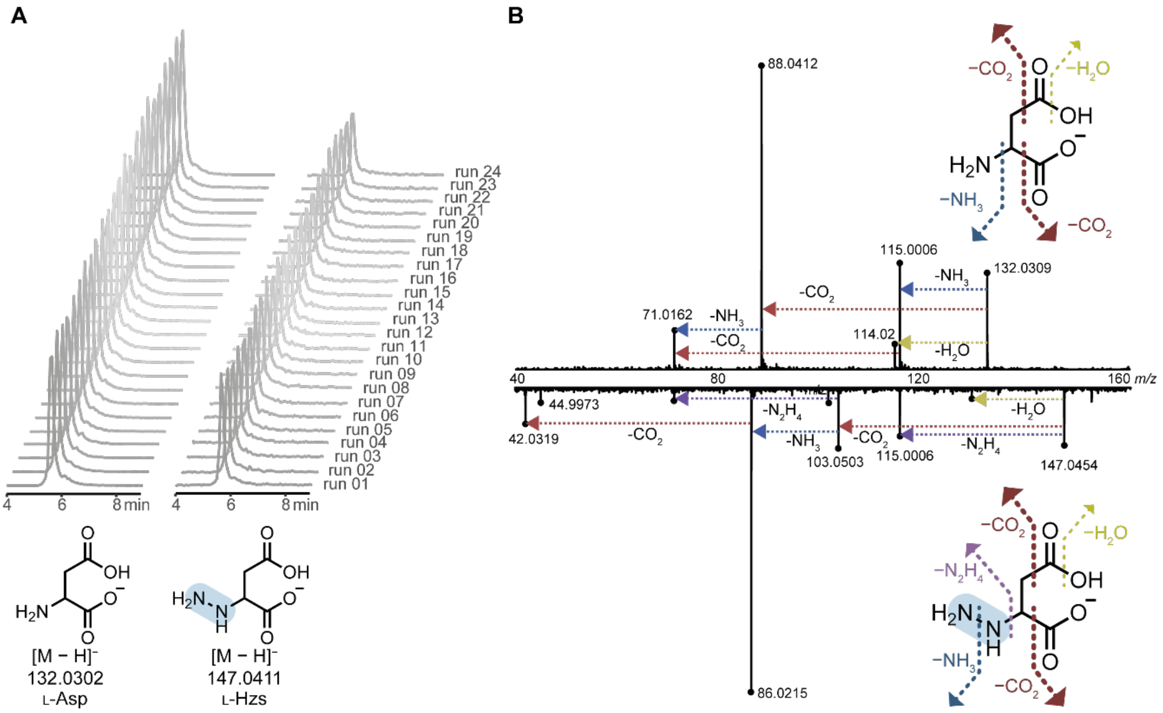


**Figure S6. L-Asp is readily distinguished from a synthetic L-Hzs standard. A)** Both underivatized L-Asp and a synthetic L-Hzs standard can be detected via LC-MS. Shown are extracted ion chromatograms for L-Asp and L-Hzs over a 12 h series of 30 minute runs. It is clear that L-hydrazinosuccinate exhibits some temperature-sensitive degradation: incubation over the timepoints indicated was associated with decreased signal, and therefore for analysis of activity assays, samples were frozen at the timepoints indicated and were run in small batches after thawing before running. **B)** Tandem mass spectrometry fragmentation patterns for L-Asp (top) and synthetic L-Hzs (fragments) show that they have both shared fragments and distinctive L-Hzs fragments, tentatively assigned to neutral losses of $CO_2$, $NH_3$, $N_2H_4$, and $H_2O$. Candidate masses detected in extracted ion chromatograms can thus be confirmed to be authentic L-Hzs via MS2 fragmentation.

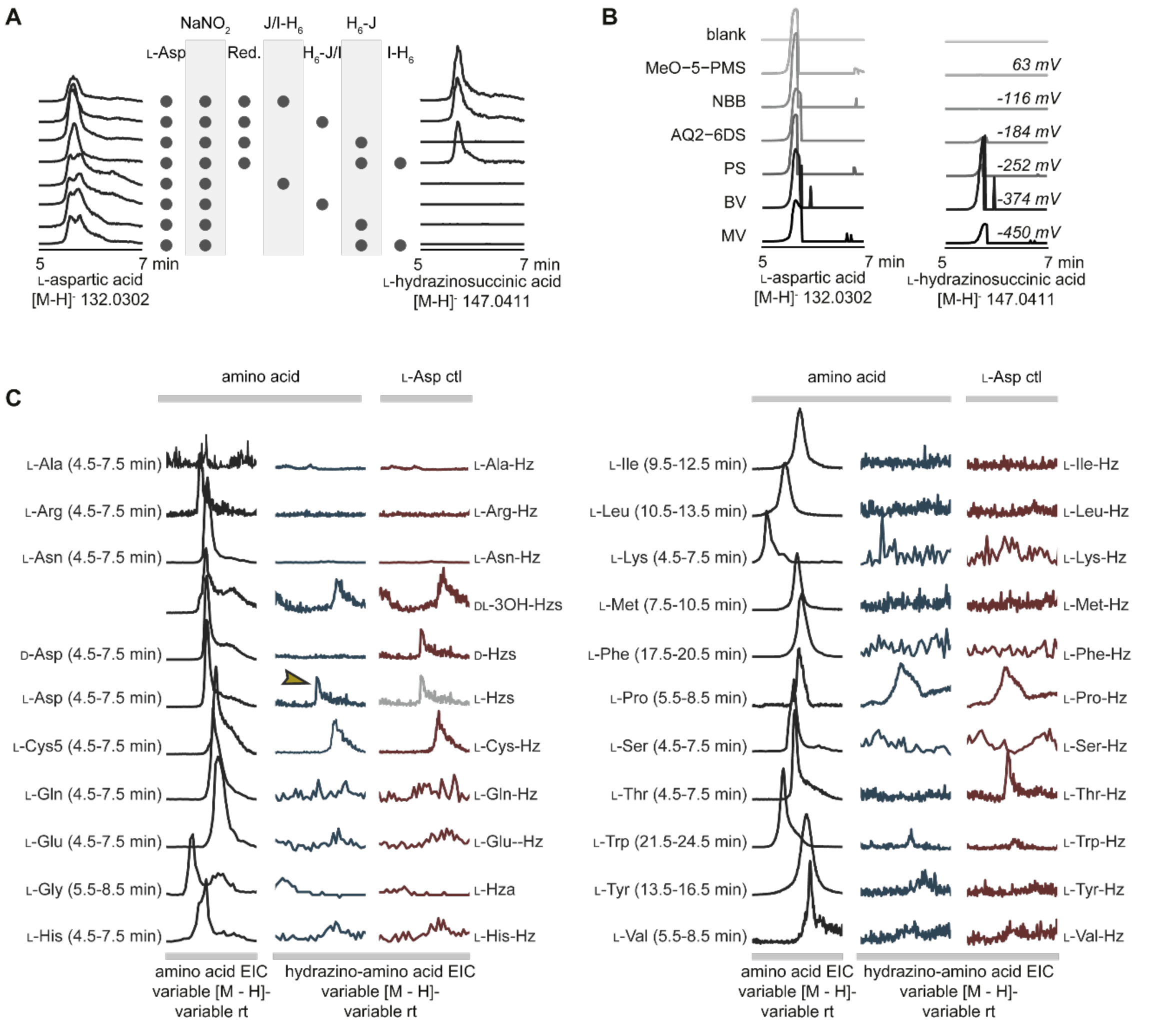

**Figure S7 Exploring KinIJ activity A)** Extracted ion chromatograms for L-Asp and L-Hzs in KinIJ activity experiments employing various combinations of $His_6$-tagged KinI and KinJ show that the tag can be present on either protein. **B)** Extracted ion chromatograms for L-Asp and L-Hzs showing KinIJ activity in the presence of various electron mediators. Little activity was observed with mediators with a redox potential over -250 mV (and maximal activity with benzyl viologen). It is not yet clear why both an electron mediator and the ferredoxin partner are required. **C)** Extracted ion chromatograms for activity assays of KinIJ with all L-amino acids (with the addition of D-Asp and DL-3OH-Asp) are shown. EICs for the unaltered amino acid are in black, and are normalized to the highest intensity. EICs for the theoretical hydrazine-amino acid product are in blue. As a control, these EICs are also shown for reactions containing L-Asp (red) – peaks present in both this control and in the sample represent background noise. The single peak seen at the L-Thr mass in the L-Asp control reflects the L-Asp +1 isotopic feature, and is thus absent in the authentic L-Thr reaction. Results in positive ion mode produced parallel results: only modification of L-Asp to form L-Hzs has been observed *in vitro* – D-Asp, DL-3OH-Asp, and L-Glu do not exhibit observed modification, and neither does Gly.

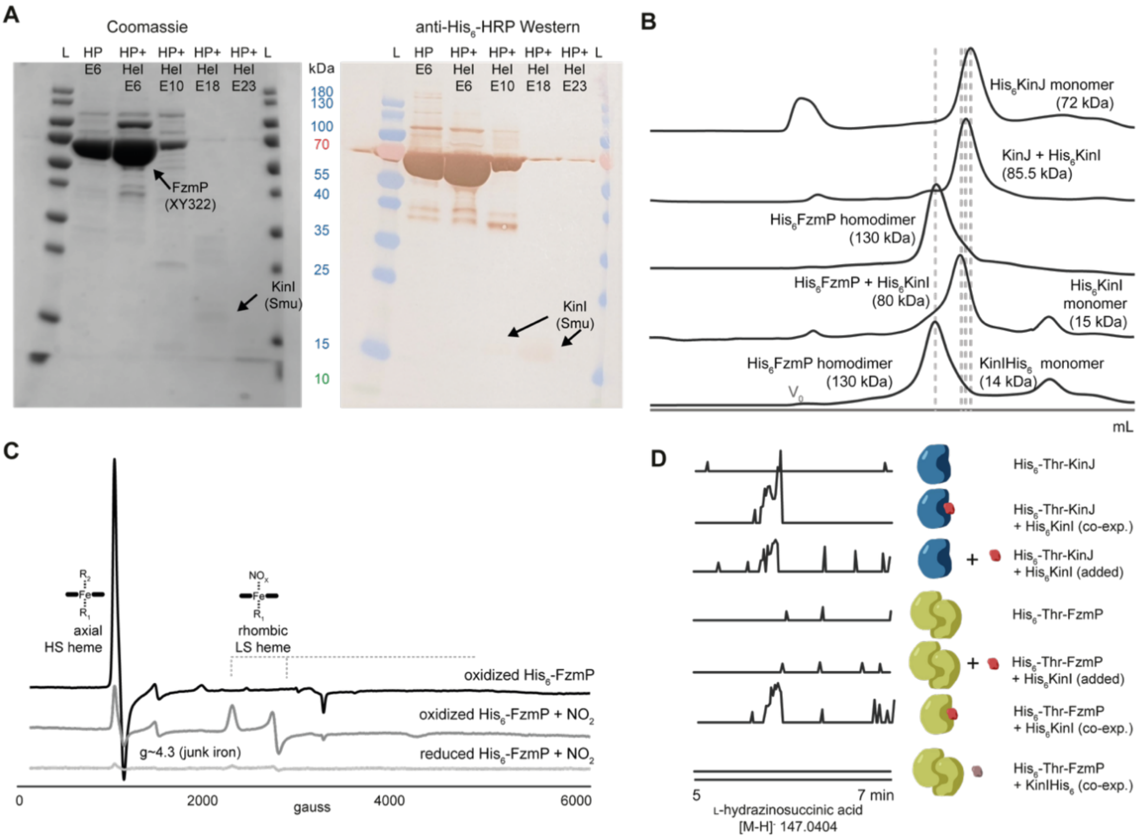

**Figure S8 Comparing KinJ and FzmP activity. A)** Expression of the KinJ homolog FzmP - SDS-PAGE & Western. **B)** Size exclusion chromatography indicates that unlike KinJ, FzmP exclusively forms a homodimer when expressed alone or in the presence of C-terminally $His_6$-tagged KinI. However, FzmP co-expressed with an N-terminally $His_6$-tagged KinI forms a heterodimer. All traces are normalized by maximum absorbance at 280 nm. **C)** X-band EPR for FzmP at 10 K supports the presence of high-spin heme species, and formation of a low-spin rhombic species after $NO_2^-$ addition. **D)** Extracted ion chromatograms for L-Hzs indicate that FzmP activity is observed with KinI as a partner, but it is observed only when FzmP is co-expressed with N-terminally tagged KinI. Without this co-expression step, the FzmP homodimer does not appear to be capable of interacting successfully with ferredoxin and thus cannot catalyze the formation of L-Hzs.

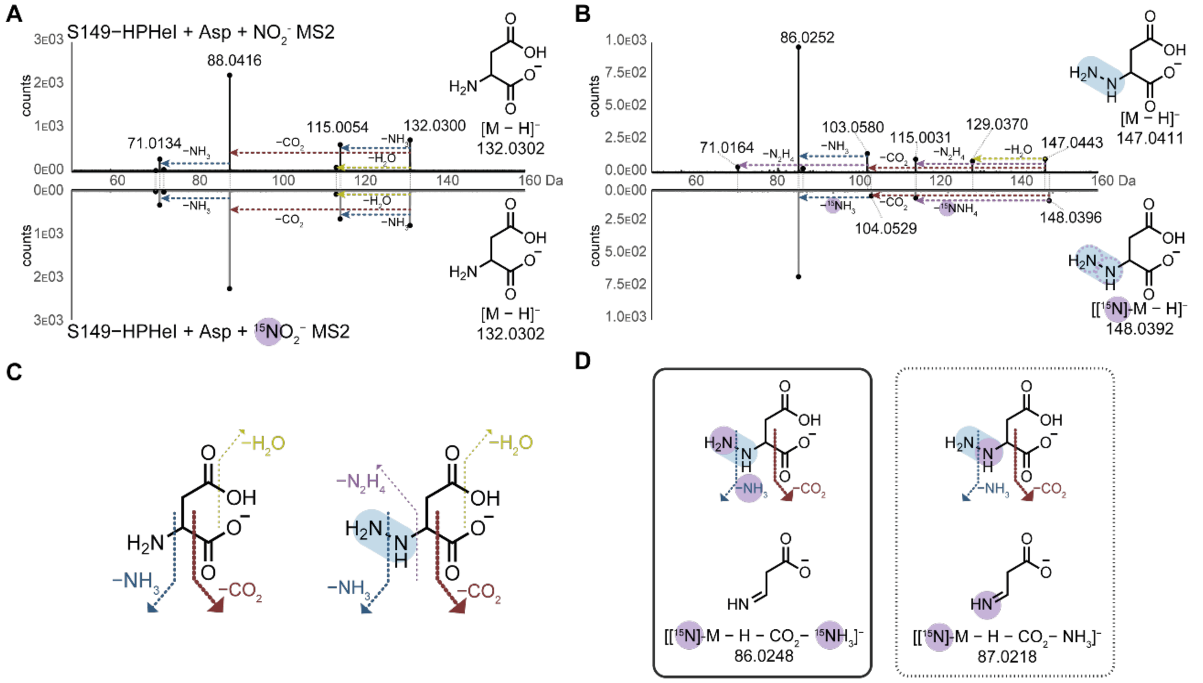


**Figure S9**. **Isotopic labeling confirms the location of nitrogen atom addition to L-Asp**. **A)** Mass spectra showing the fragmentation of L-Asp (132.0302 *m/z*) in an FzmP reaction carried out using either $NO_2^-$ (top) or $^{15}NO_2^-$ (bottom). **B)** Spectra showing the fragmentation of L-Hzs (top: 147.0411 *m/z*) and $^{15}$N- L-Hzs (bottom: 148.0392 *m/z*) after the FzmP reaction with $NO_2^-$ and $^{15}NO_2^-$ respectively. **C)** Proposed neutral losses responsible for the generation of the observed MS2 fragment ions. **D)** The most abundant fragment ion for L-Hzs is consistent with the loss of a $CO_2$ and an $NH_3$ group. Retention of the $^{15}$N atom in that fragment would reflect its probable proximal position; however, this isotopically enriched fragment is not observed. Fragmentation of the enzymatically produced $^{15}$N-labeled L-Hzs is consistent with the loss of a labeled distal nitrogen, confirming location of nitrogen installation. $^{15}$N labeling also supports the proposed assignments of other fragment ions: the 103 Da *m/z* species – representing a neutral $CO_2$ loss – is replaced by a 104 Da *m/z* species for $^{15}$N- L-Hzs, while the 115 Da *m/z* species – representing $N_2H_4$ loss, and thus loss of the isotopically labeled nitrogen – is not.

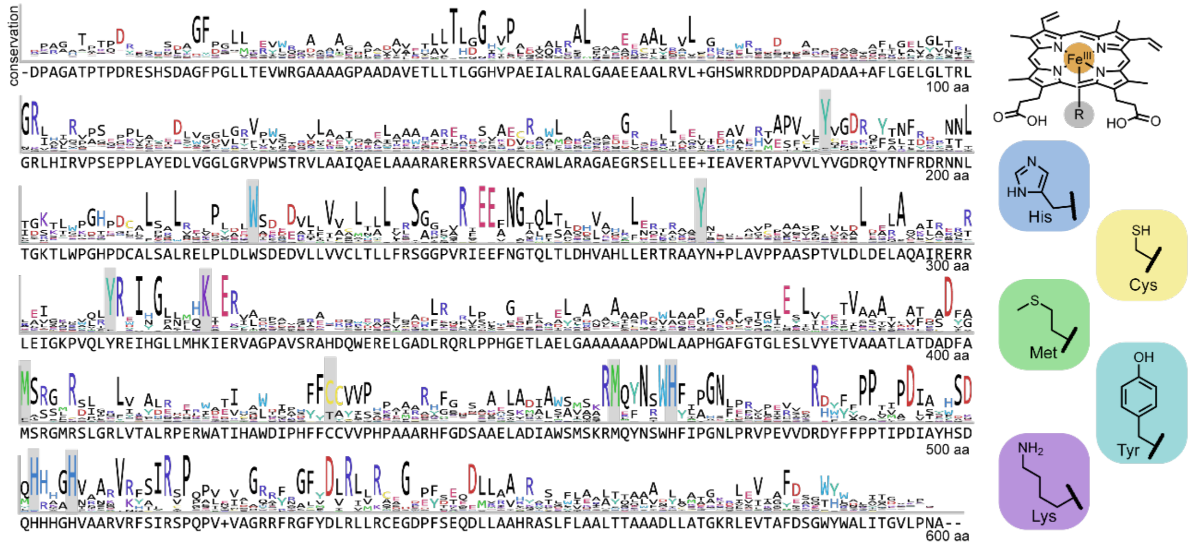


**Figure S10. Prioritizing KinJ variants.** A MAFFT multiple sequence alignment made using the representative node sequences yields information about conserved residues. Residues with the ability to bind metals (C, D, E, H, M) and hemes (C, H, M, K, Y)[24] or to transport electrons (W, Y) are shown in color, and residues altered via mutagenesis in this work are highlighted.

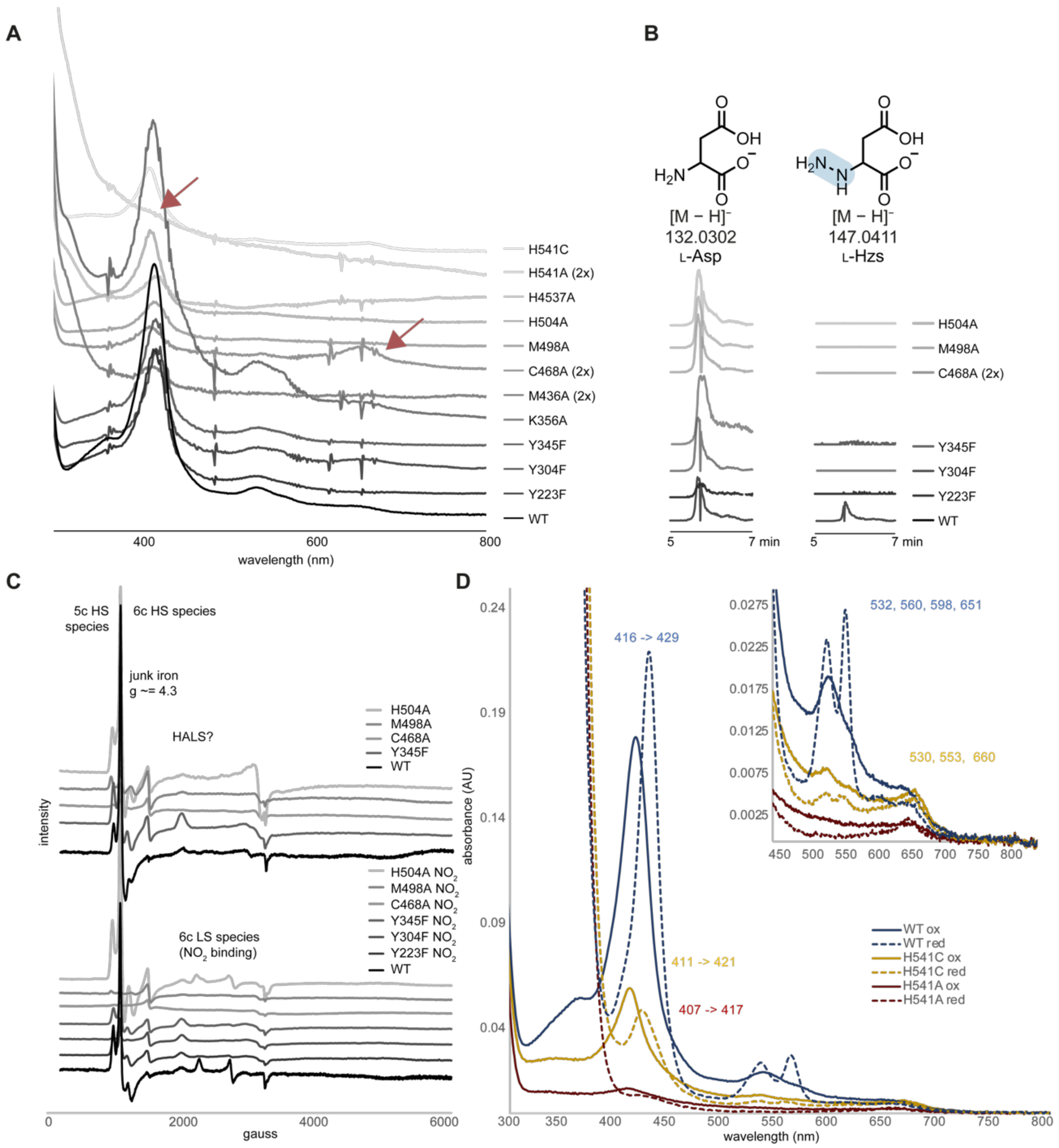


**Figure S11. Investigation of KinJ variants. A**) UV-vis spectroscopy of KinJ variants. Heme levels varied by prep, but significant decreases in heme loading or alteration of spectral features were observed only for a subset of variants. Heme binding was absent in H541A variant, while heme-associated spectral features (Soret and Q-band peaks) were visibly altered in M498A and C468A. These two variants had an additional spectral feature at ~650 nm and a significantly decreased Soret peak. **B**) Substitution of highly conserved residues eliminated N–N bond formation. **C**) EPR analysis of select variants shows the same mix of 5- and 6-coordinate high-spin species in the absence of $NO_2^-$and the $NO_2^-$-bound 6-coordinate low spin species (though the amplitude of this low-spin feature is significantly decreased.) **D**) Alanine replacement of the proposed axial ligand (H541A) results in loss of heme binding. Cysteine replacement (H541C), however, yields a construct with an alternate axial ligand and heme binding is recovered. Compared to the WT species, this heme exhibits spectral characteristics consistent with a high-spin thiolate heme (previously reported in myoglobin H93C variants),[25] including a broadened and blue-shifted Soret peak and a comparatively strong absorption feature above 650 nm.[26] This supports the proposal that loss of heme binding is due exclusively to loss of the axial ligand and not other factors (loss of H-bonding, loss of an aromatic residue, loss of steric bulk) that might alter the active site environment.

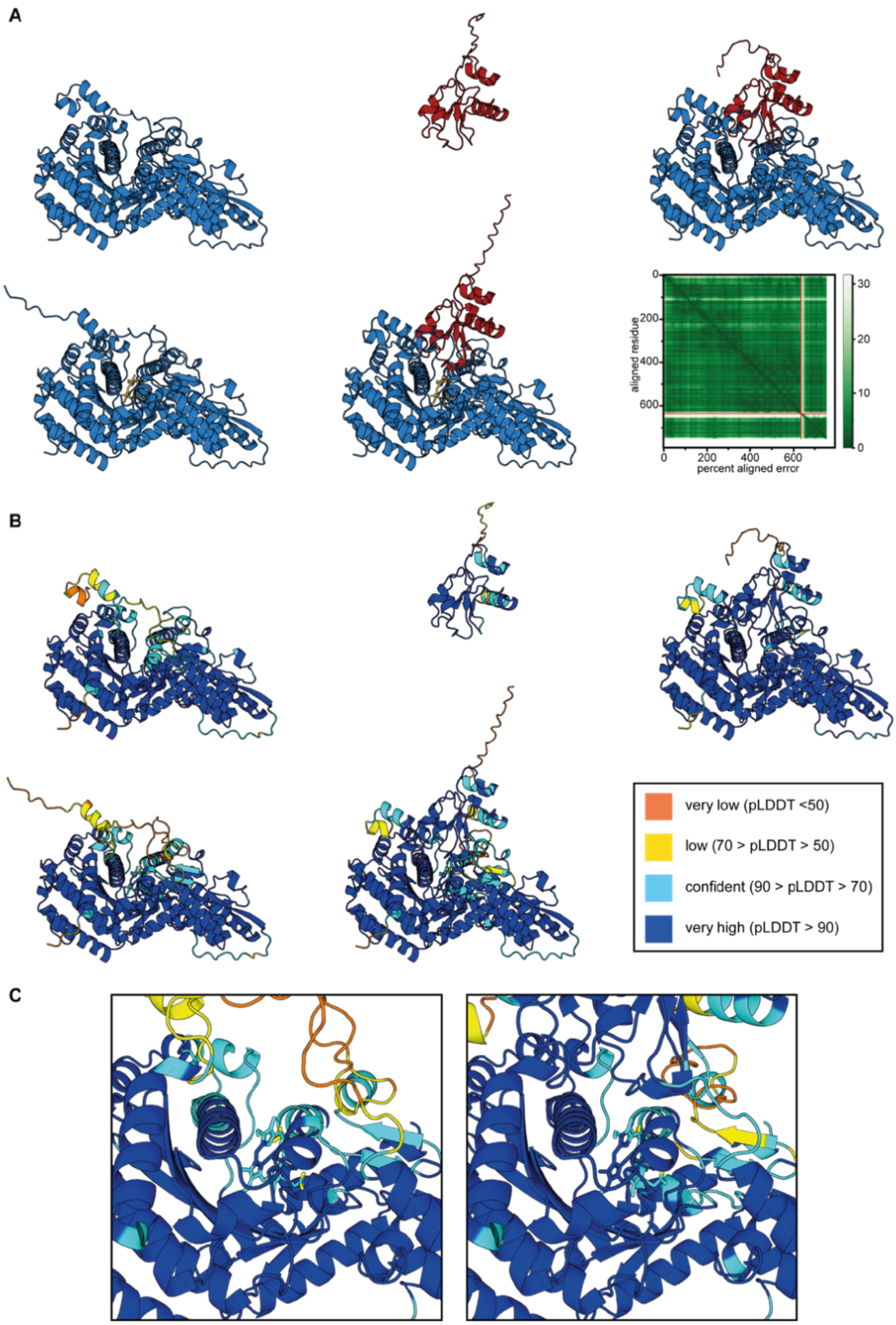


**Figure S12. The predicted fold for KinJ is consistent across models and oligomeric states. A**) Predicted structures were generated for KinJ (top left, blue) and KinI (top center, red) via AlphaFold2. Folds for both proteins are preserved in the the KinIJ heterodimer, modeled via AlphaFold2 Multimer (top right). Predicted folds for KinJ and KinIJ were also produced with a heme cofactor via AlphaFold3 (bottom left and center.) The predicted fold for KinJ is consistent across these models. The percent aligned error for the KinIJ heterodimeric complex (bottom right) strongly supports multimer formation. Lower PAE values (green) are better; PAE scoring is as high between KinI and KinJ as it is within either structure, supporting heterodimer formation. **B**) The pLDDT scoring is shown for KinJ (top left), KinI (top center) and KinIJ (top right) produced by AlphaFold2/AlphaFold Multimer, and holo KinJ/KinIJ modeled by AlphaFold3 (bottom left, bottom center). The N-termini of both KinI and KinJ are lower-confidence, but modeling KinIJ as a heterodimer results in an interaction between the KinJ N-terminus and the ferredoxin that stabilizes much of that region of KinJ. **C**) The proposed active site pocket (left) is also higher-confidence when modeled with KinI present (right).

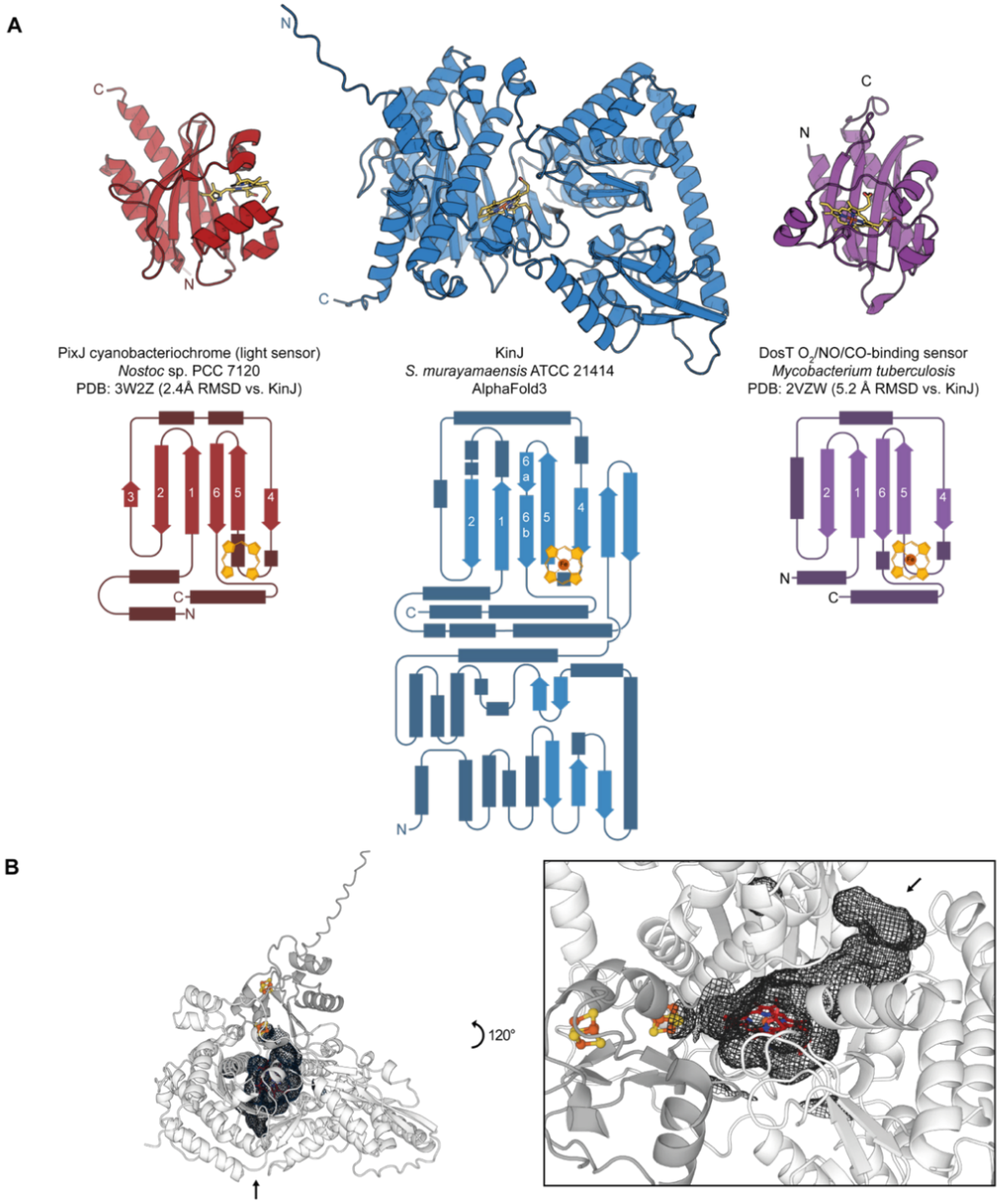


**Figure S13. The C-terminal KinJ domain has unexpected structural homology with GAF domains in sensor proteins. A**) The C-terminal domain of the predicted KinJ (blue) structure is aligned with the structures of PixJ (red) and DosT (purple),[27–29] chosen as representative cofactor-bound GAF domains identified via DALI. The GAF domain of PixJ contains a bilin cofactor and constitutes the light-sensing domain in a cyanobacteriochrome. DosT is a histidine kinase hypoxia sensor, and the GAF domain of DosT contains a heme cofactor, which binds $O_2$, NO, and CO. The C-terminal domain of KinJ has a structurally homologous core. Similar to the heme-binding GAF domain, KinJ lacks the 3rd beta strand and instead contains an alpha helix. Compared to both bilin- and heme-binding GAF domains, the KinJ C-terminal domain is more complex, containing an extra pair of beta-strands, a bifurcated final beta strand, and several additional helices, along with an N-terminal domain with no identifiable homology to known or predicted protein folds. The axial ligand for DosT and the proposed axial ligand for KinJ are in a similar loop region. While a bilin cofactor does not have an axial ligand, it binds in a similar area of the GAF domain fold. **B**) KinJ contains a central cavity (left); this cavity is large enough to contain the heme that is modeled in by AlphaFold3 and bound to the axial ligand H541 (right). (In the pictured models, the 4Fe-4S clusters are positioned via alignment of KinI against PDB: 3EUN, a 2[4Fe-4S] ferredoxin[22]). Additional space remains open on the proximal side of the heme cofactor, consistent with a potential active site where two substrates must access the heme cofactor. A potential tunnel for substrate access is present (arrow points to entrance).

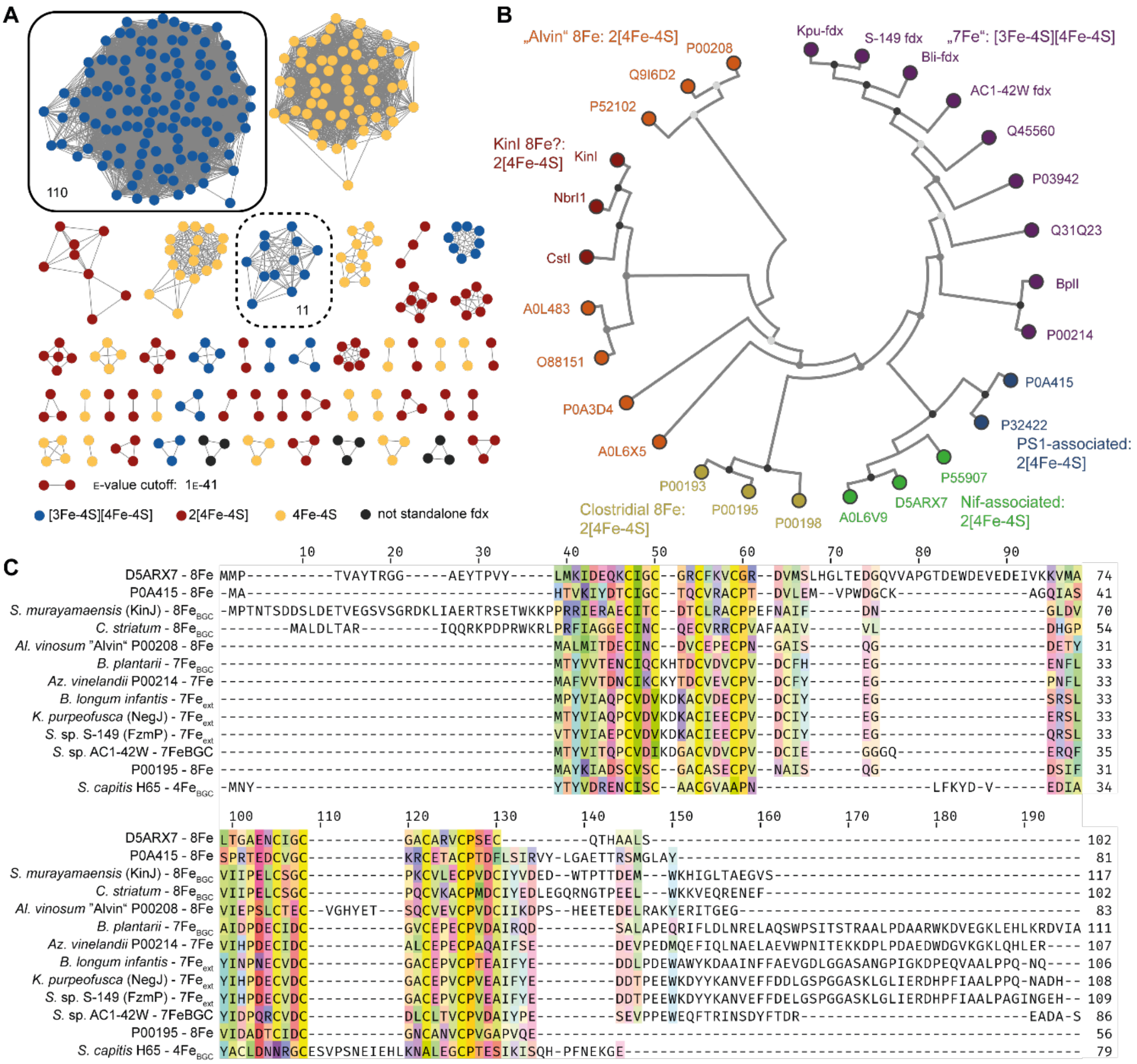


**Figure S14. Partner ferredoxins are not all homologous to KinI. A**) Lack of a ferredoxin in the genomic neighborhood of FzmP delayed identification of the requirement for a ferredoxin, and other KinJ homologs lack a clear partner (**Fig. 2B**). Using the representative KinJ homologs from **Fig. 1**, homologs from genomic neighborhoods lacking a ferredoxin were identified. For homologs present in IMG genomes, the full genomes were searched for all genes encoding proteins with 4Fe-4S motifs using all extant 4Fe-4S ferredoxin Pfams. Protein sequences longer than 150 amino acids were removed, and a sequence similarity network was constructed for the remaining ferredoxins with a cutoff at 1E-41. In these genomes, 110/149 species encode a specific subtype of 7Fe ferredoxin with a homolog found in a subset of KinJ-associated BGCs; 11 encode a slightly divergent 7Fe ferredoxin. We identified those 7Fe ferredoxins as the most likely candidates for a non-BGC-encoded partner – and support from a recent publication identifying a KinJ homolog involved in negamycin biosynthesis suggests that this is correct.[30] Monocluster 4Fe-4S candidates were relatively widespread but no subgroups were as universally distributed. **B**) Representative KinJ partner ferredoxins (including members of the 7Fe subclass) were aligned via MAFFT against previously characterized ferredoxins belonging to characterized 4Fe-4S ferredoxin subclasses and visualized in an unrooted maximum likelihood tree generated in MEGA. Black circles over nodes represent bootstrap values >75%; grey circles represent bootstrap values >50%; light grey circles represent bootstrap values under 50%. Notably, KinJ-associated ferredoxins are heterogeneous, falling into several groups. KinIs form a subgroup distinct from other 8Fe ferredoxins such as "Alvin" ferredoxin from *Allochromatium vinosum*.

The 7Fe ferredoxins are related to the 7Fe ferredoxin from *Azotobacter vinelandii*, though some are encoded in BGCs containing KinJ homologs and some are encoded elsewhere in the genome. Not shown in the tree are ferredoxins from *Staphylococcus* BGCs: these encode a mononuclear 4Fe-4S ferredoxin, and their genomes lack close homologs of both the 8Fe and 7Fe ferredoxins. **C)** Sequence alignment for members of each class. Compared to the "Alvin" ferredoxins, KinI homologs have an extended N-terminus prior to the first Cx2Cx2Cx3CP motif and lack the insertion in the second $Cx_2Cx_6CX_3CPx_2C$ motif seen in the Alvin ferredoxin. The 7Fe ferredoxins have an initial $Cx_2(C/V)x_4CX_3CPx_2C$ motif (the second cysteine does not coordinate an iron atom in 7Fe ferredoxins), a second $Cx_2Cx_2Cx_3CP$ motif and an extended C-terminus. The *Staphylococcus* ferredoxin ScuI clearly lacks cysteines required for dicluster binding.

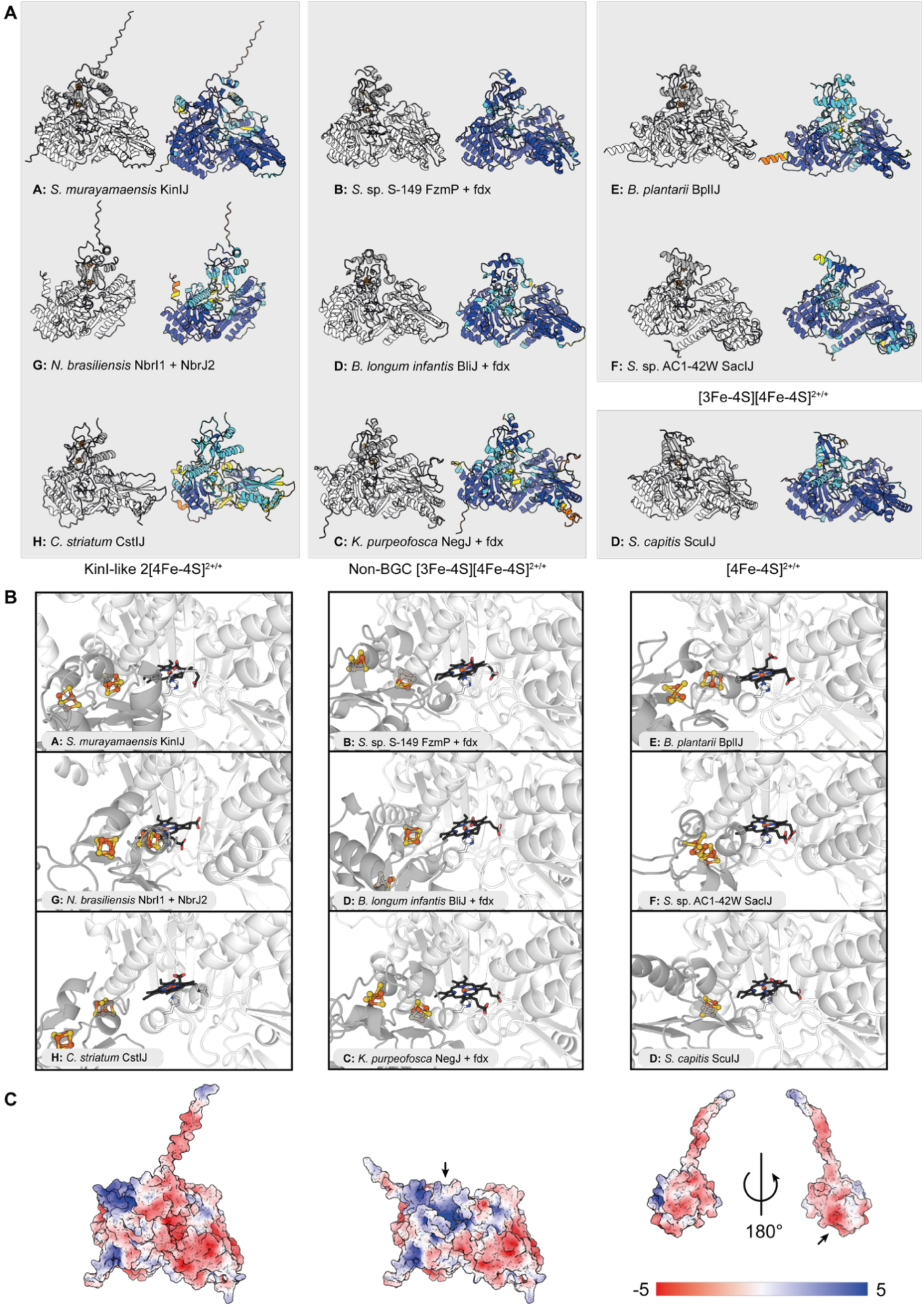


**Figure S15. Predicted structures for KinJ homologs and their partner ferredoxins. A)** AlphaFold3 predictions for all KinJ homologs share the same core fold, with pLDDT (shown to the right of each model) varying primarily at the N- and C- termini. All ferredoxin partners were identified are shown with iron-sulfur clusters modeled in via alignment against the closest structural relative (PDB: 3EUN for KinI-like 8Fe ferredoxins, PDB: 7FDR for *Az.*

*vinelandii*-like 7Fe ferredoxins, and PDB: 1IQZ for the ScuI 4Fe ferredoxin.) Grouping by KinJ sequence similarity does not map onto ferredoxin family or onto genome neighborhood type. *N. brasiliensis* has two BGCs utilizing KinJ homologs; one possesses a KinI-like ferredoxin and the other has a 4Fe-4S ferredoxin in the active site. However, AlphaFold3 confidence metrics provide modestly stronger support for the binding of the KinI ferredoxin found in the other BGC. Differences in KinJ homolog structures are visible: members of groups G and H, for example, have a comparatively truncated N-terminal domain. **B)** Ferredoxins are shown in darker gray and KinJ homologs in white. As the differences in FeS site positioning and the positioning of the key 4 beta sheets in the ferredoxin fold indicate, the orientations of some ferredoxin partners vary, particularly those for which the pLDDT scoring near the heme protein/ferredoxin interface was low. However, the nearest ferredoxin FeS cluster is consistently modeled in close proximity to the heme-binding site, and the active site pocket is not significantly affected by changes at the N- or C-termini of homologs. **C)** An acidic patch on the model of KinI is positioned well to bind a basic region surrounding the tunnel into the KinJ active site. Left: KinIJ complex. Center: KinJ, with arrow pointing into basic cleft surrounding the tunnel to the active site. Right: KinI, viewed both in the orientation seen at in the KinIJ complex and rotated 180° to reveal the acidic patch.

**Supplementary Tables**

**Supplementary Table S1: Strains used in this study**

| **Strain** | **Source** |
|---|---|
| *Escherichia coli* LOBSTR (DE3) | Kerafast |
| *Escherichia coli* Top10 | Invitrogen |
| *Escherichia coli* NEB-5α | NEB |
| *Escherichia coli* “E cloni” 10G | Lucigen |
| *Streptomyces murayamaensis* ATCC 21414 | ATCC |
| *Streptomyces* sp. NRRL S-149 | Metcalf and van der Donk groups |
| *Streptomyces* sp. XY322 | Metcalf and van der Donk groups |

**Supplementary Table S2:** Primers used during the assembly of non-codon-optimized

| Construct(s) | Forward Seq | Reverse Seq | PCR Template | Vector | Method |
|---|---|---|---|---|---|
| Smu HrJ (WT) | ATGCATCATATGACAAGTTCCGCCGAG | ATGCATCTCGAGTCACGAGACCTGCTGCG | *S. murayamaensis* ATCC 21414 gDNA | pET-28a | Smu KinJ into pET-28a via NdeI/XhoI |
| S-149 HrP | AGCGGCCTGGTGCCGCGCGGCAGCCATATGATGCTTCTGAAGACGC | CCTTTCGGGCTTTGTTAGCAGCCGGATCCTCGAGTCAGGAGACATGGAAG | S. sp. S-149 gDNA | pET-15b | S sp S-149 into pET-15b via Gibson, vector digest |
| S-149 P (MCS1) and Smu HeI (MCS2) in pCDFDuet-1 | ATAAGGAGATATACCATGCTTCTGAAGACGCCCG | GCATTATGCGGCCGCTCAGGAGACATGGAAGTCCCG | S-149 His6-Thr-FzmP in pET15b | pCDFDuet-1 -/HeI | S. sp. S-149 FzmP into MCS1 of pETCDFDuet vectors |
| See above | GCGGCCGCATAATGCTTAAGT | GGTATATCTCCTTATTAAAGTTAAA | pCDFDuet-1 -/HeI | pCDFDuet-1 -/HeI | Linearized vector for FzmP insertion into MCS1 (Gibson) |
| XY322 HrP | AGCGGCCTGGTGCCGCGCGGCAGCCATATGATGCAGCTGAAGACGCCC | CCTTTCGGGCTTTGTTAGCAGCCGGATCCTCGAGTCAGGAGCCATGGAAGGC | S. sp. XY-322 gDNA | pET-15b | S sp XYX332 FzmP into pET-15b via Gibson, vector digest |
| Smu KinI in MCS2 for pCDFDuet-1, untagged (-/I) | ATGCTACATATGCCCACGAACACTTCG | ATGCTACTCGAGTCATGACACACCCTCAGC | *S. murayamaensis* (Smu) gDNA | pCDFDuet-1 | Cloning of Smu-KinI into Duet vector MCS2 via NdeI/XhoI digest |
| Smu -/HeI | ATCACCACGATGACGACGACAAGATGCCCACGAACACTTC | GATGGTGATGGCTGCTGCCCATATGTATATCTCCTTCTTATACTTAAC | Smu -/I | pCDFDuet-1 -/I | Insertion of His6-Ek at the N-terminus of KinI in MCS2 in Duet vectors |
| Smu KinI-His$_5$ (IH$_5$) in pCDFDuet-1 MCS2 | ACCACCATCATTAATAACTCGAGTCTGGTAAAGAAACCG | GGTGGCTCGCGCCGATTGACACACCCTCAGCGG | Smu -/I | pCDFDuet-1 -/I | Insertion of His5 at the C-terminus of KinI in MCS2 in Duet vectors |
| Smu KinI-His$_6$ (IH) in pCDFDuet-1 MCS2 | GCCACCACCACCATCATCACTAATAACTCGAGTCTGG | CCAGACTCGAGTTATTAGTGATGATGGTGGTGGTGGC | Smu -/IH5 | pCDFDuet-1 -/IH5 | Adding C-terminal His to KinI-His5 via Quikchange mutagenesis, yielding KinI-His6 |
| Smu HJHeI, HJIH | ATGCTAGGATCCTATGACAAGTTCCGCCGAG | ATGCTAAAGCTTTCACGAGACCTGCTGCG | Smu HrJ | pCDFDuet-1 -/HeI, pCDFDuet-1 -/IH, pCDFDuet-1 -/I | Cloning of Smu KinJ into Duet vectors at MCS1 (retaining the vector N-terminal His6 tag) via BamHI/HindIII digest. |
| Smu HeJI | ATGCTAGGATCCTGATGACGACGACAAGATGACAAGTTCCGCCGAG | ATGCTAAAGCTTTCACGAGACCTGCTGCG | Smu HrJ | pCDFDuet-1 -/I | Addition of Ek-KinJ into MCS1 in Duet vectors (His6 tag from the vector maintained) via BamHI/HindIII digest |
| Smu JHeI, JIH, JI | CATCACCACCATCATTGAAAGCTTGCGGCCGC | GTGAGATGCACCCGAGACCTGCTGCGGC | Smu HJ/HeI, Smu HJ/IH, Smu HJ/I | pCDFDuet-1 HJ/IH, pCDFDuet-1 HJ/HeI, pCDFDuet-1 HJ/I | Removal of N-terminal His6 for KinJ via around-the-horn PCR and ligation |
| Smu JHI | CATCACCACCATCATTGAAAGCTTGCGGCCGCATAATGC | GTGAGATGCACCCGAGACCTGCTGCGGCTGC | Smu J/I | pCDFDuet-1 -/I | Addition of C-terminal His6 for KinJ via around-the-horn PCR and ligation |
| Smu HrJ Y223F | CGCGGAAGTTGGTGAAGAGCTTCCCCTCC | GGAGGGGAAGCTCTTCACCAACTTCCGCG | Smu His6-Thr-KinJ in pET-28a | Smu HrJ (pET-28a) | SmuKinJ QuikChange mutagenesis: Y233F |
| Smu HrJ Y304F | GGGACCGCGTTGAAGTTGCCGCGGG | CCCGCGGCAACTTCAACGCGGTCCC | Smu His6-Thr-KinJ in pET-28a | Smu HrJ (pET-28a) | SmuKinJ QuikChange mutagenesis: Y304F |
| Smu HrJ C468S | GGCACCACGCAGCTGAAGAAGTCGGTG | CACCGACTTCTTCAGCTGCGTGGTGCC | Smu His6-Thr-KinJ in pET-28a | Smu HrJ (pET-28a) | SmuKinJ QuikChange mutagenesis: C468S |
| Smu HrJ M498A | CAGGAGTTGTACTGCGCGCGGGACGACATCGC | GCGATGTCGTCCCGCGCGCAGTACAACTCCTG | Smu His6-Thr-KinJ in pET-28a | Smu HrJ (pET-28a) | SmuKinJ QuikChange mutagenesis: M498A |
| Smu HrJ H504A | GTTCCCGGCGATGAAGGCCCAGGAGTTGTACTGC | GCAGTACAACTCCTGGGCCTTCATCGCCGGGAAC | Smu His6-Thr-KinJ in pET-28a | Smu HrJ (pET-28a) | SmuKinJ QuikChange mutagenesis: H504A |

**Supplementary Table S3: List of KinI, KinJ/FzmP, and co-expression constructs used for heterologous protein expression**

| Name | Backbone | MCS1 | MCS2 (if applicable) | Species |
|---|---|---|---|---|
| Smu HrJ | pET-28a | $His_6$-Thr-KinJ | | *Streptomyces murayamaensis* ATCC 21414 |
| Smu HrJ Y223F | pET-28a | $His_6$-Thr-KinJ | | *Streptomyces murayamaensis* ATCC 21414 |
| Smu HrJ Y304F | pET-28a | $His_6$-Thr-KinJ | | *Streptomyces murayamaensis* ATCC 21414 |
| Smu HrJ Y346F | pET-28a | $His_6$-Thr-KinJ | | *Streptomyces murayamaensis* ATCC 21414 |
| Smu HrJ K356A | pET-28a | $His_6$-Thr-KinJ | | *Streptomyces murayamaensis* ATCC 21414 |
| Smu HrJ M436A | pET-28a | $His_6$-Thr-KinJ | | *Streptomyces murayamaensis* ATCC 21414 |
| Smu HrJ C468S | pET-28a | $His_6$-Thr-KinJ | | *Streptomyces murayamaensis* ATCC 21414 |
| Smu HrJ M498A | pET-28a | $His_6$-Thr-KinJ | | *Streptomyces murayamaensis* ATCC 21414 |
| Smu HrJ H504A | pET-28a | $His_6$-Thr-KinJ | | *Streptomyces murayamaensis* ATCC 21414 |
| Smu HrJ H537A | pET-28a | $His_6$-Thr-KinJ | | *Streptomyces murayamaensis* ATCC 21414 |
| Smu HrJ H541A | pET-28a | $His_6$-Thr-KinJ | | *Streptomyces murayamaensis* ATCC 21414 |
| Smu HrJ H541C | pET-28a | $His_6$-Thr-KinJ | | *Streptomyces murayamaensis* ATCC 21414 |
| Smu IH | pCDFDuet-1 | | KinI-$His_6$ | *Streptomyces murayamaensis* ATCC 21414 |
| Smu HeI | pCDFDuet-1 | | $His_6$-Ek-KinI | *Streptomyces murayamaensis* ATCC 21414 |
| Smu HeJI | pCDFDuet-1 | $His_6$-Ek-KinJ | KinI | *Streptomyces murayamaensis* ATCC 21414 |
| Smu JHeI | pCDFDuet-1 | KinJ | $His_6$-Ek-KinI | *Streptomyces murayamaensis* ATCC 21414 |
| Smu JIH | pCDFDuet-1 | KinJ | KinI-$His_6$ | *Streptomyces murayamaensis* ATCC 21414 |
| Smu JHI | pCDFDuet-1 | KinJ-$His_6$ | KinI | *Streptomyces murayamaensis* ATCC 21414 |
| S-149 HrP | pET-15b | $His_6$-Thr-FzmP | | *Streptomyces* sp. S-149 |
| XY332 HrP | pET-15b | $His_6$-Thr-FzmP | | *Streptomyces* sp. XY332 |
| S-149 PHeI | pCDFDuet-1 | FzmP | His6-Ek-KinI | *Streptomyces* sp. NRRL S-149 & *Streptomyces murayamaensis* ATCC 21414 |

**Supplementary Table S4: DALI results for KinJ**

Results from the full PDB are dominated by a large number of crystal structures for a specific GAF-domain-containing histidine kinase sensor from *M. tuberculosis*.

| Chain | Z-score | RMSD | LALI | NRes | %ID | Protein | PDB description |
|---|---|---|---|---|---|---|---|
| 5DFX-A | 9.7 | 3.1 | 137 | 158 | 6 | Slr1393 | Chromophore-bound histidine kinase (GAF3 domain from Slr1393) |
| 2Y79-A | 9.7 | 2.7 | 124 | 150 | 13 | DosS | E87A mutant of GAF domain from the *M. tuberculosis* redox sensor DosS, a histidine kinase |
| 2Y8H-B | 9.7 | 2.7 | 127 | 148 | 13 | DosS | E87G mutant of GAF domain from *M. tuberculosis* DosS |
| 2W3G-B | 9.6 | 2.7 | 124 | 145 | 13 | DosS | Air-oxidized GAF domain from *M. tuberculosis* DosS |
| 2W3F-B | 9.6 | 2.8 | 124 | 144 | 14 | DosS | Reduced GAF domain from *M. tuberculosis* DosS |
| 2W3G-A | 9.6 | 2.7 | 123 | 150 | 13 | DosS | Air-oxidized GAF domain from *M. tuberculosis* DosS |
| 3VV4-B | 9.6 | 2.7 | 137 | 160 | 11 | TePixJ | GAF domain from the cyanobacteriochrome TePixJ |
| 2W3E-B | 9.6 | 2.7 | 124 | 145 | 13 | DosS | Oxidized GAF domain from *M. tuberculosis* DosS |
| 3W2Z-A | 9.6 | 3.3 | 148 | 178 | 7 | AnPixJ | GAF domain from the cyanobacteriochrome AnPixJ |
| 2Y79-B | 9.6 | 2.7 | 124 | 146 | 14 | DosS | E87 mutant of GAF domain from *M. tuberculosis* DosS |

Results from the PDB90 are more diverse, with a range of GAF domains containing heme or cyanobacteriochrome cofactors. While the level of similarity with any of these structures is quite low, matches are consistently for cyanobacteriochrome and heme-binding GAF domains.

| Chain | Z-score | RMSD | LALI | NRes | %ID | Protein | PDB description |
|---|---|---|---|---|---|---|---|
| 3W2Z-A | 9.6 | 3.3 | 148 | 178 | 7 | AnPixJ | GAF domain from the cyanobacteriochrome AnPixJ |
| 6OAP-A | 9.5 | 10.0 | 164 | 305 | 8 | PPHK | Dual sensor histidine kinase in the green light-absorbing Pg state |
| 3CIT-A | 99.5 | 2.8 | 133 | 155 | 8 | | GAF domain from a putative histidine kinase in *P. syringae* pv. tomato |
| 2VZW-B | 9.4 | 3.0 | 124 | 149 | 15 | DosT | The heme-bound GAF domain of the sensory histidine kinase DosT of *M. tuberculosis* |
| 2Y8H-A | 9.4 | 2.9 | 121 | 152 | 14 | DosS | E87G mutant of GAF domain from *M. tuberculosis* DosS |
| 6MGH-D | 9.3 | 3.0 | 135 | 152 | 8 | miRFP670nano | Monomeric near-infrared fluorescent protein miRFP670nano (evolved from cyanobacteriochrome) |

| | | | | | | | |
|---|---|---|---|---|---|---|---|
| 5M85-A | 9.3 | 2.8 | 130 | 159 | 7 | Slr1393 | The intermediate state of GAF3 domain from Slr1393 (*Synechocystis* sp. PCC6803) |
| 4FOF-A | 9.3 | 3.4 | 141 | 168 | 7 | PixJ | The blue-light absorbing form of the *T. elongatus* PixJ GAF domain |
| 3HCY-A | 9.2 | 2.8 | 128 | 145 | 11 | | Putative two-component sensor histidine kinase protein from *S. meliloti* 1021 |
| 6XHH-B | 9.1 | 3.3 | 141 | 180 | 4 | JSC1_58120g3 | Far-red absorbing dark state of JSC1_58120g3 with bound 18-1,18-2 dihydrobiliverdin IXa (DHBV), the native chromophore precursor |

30 M. Wang, Z. W. Wei and K. S. Ryan, *Nat Chem Biol*, DOI:10.1038/s41589-025-01898-0.
